## Supplemental Materials for "Does slow oscillation-spindle coupling contribute to sleep-dependent memory consolidation? A Bayesian meta-analysis"

#### Contents


**1 PRISMA statements checklist** **1**

**2 Study characteristics and effect size** **4**

**3 Risk of Bias Assessment** **4**

**4 Model Diagnostics** **8**

**5 Effect size-level forest plot** **10**

**6 Pareto *K* diagnosis statistics** **14**

**7 Moderator models** **18**

**8 Posterior distribution of moderators** **19**

**9 Interaction models and sensitivity analysis** **23**

**10 Frequentist analysis** **27**

**11 Funnel plots for publication bias** **31**

**12 Simulation of circular-linear correlation and standardization** **32**

**13 SO-slow SP spatial-temporal analysis** **35**

#### PRISMA Statements Checklist

| Section and Topic | Item # | Checklist item | Location where item is reported |
|---|---|---|---|
| **TITLE** | | | |
| Title | 1 | Identify the report as a systematic review. | Title |
| **ABSTRACT** | | | |
| Abstract | 2 | See the PRISMA 2020 for Abstracts checklist. | ☑ |
| **INTRODUCTION** | | | |
| Rationale | 3 | Describe the rationale for the review in the context of existing knowledge. | Introduction |
| Objectives | 4 | Provide an explicit statement of the objective(s) or question(s) the review addresses. | Introduction |
| **METHODS** | | | |
| Eligibility criteria | 5 | Specify the inclusion and exclusion criteria for the review and how studies were grouped for the syntheses. | Method |
| Information sources | 6 | Specify all databases, registers, websites, organisations, reference lists and other sources searched or consulted to identify studies. Specify the date when each source was last searched or consulted. | Method |
| Search strategy | 7 | Present the full search strategies for all databases, registers and websites, including any filters and limits used. | Method |
| Selection process | 8 | Specify the methods used to decide whether a study met the inclusion criteria of the review,, and if applicable, details of automation tools used in the process. | Method |
| Data collection process | 9 | Describe method of data extraction from reports (e.g., piloted forms, independently, in duplicate) and any processes for obtaining and confirming data from investigators. | Method |
| Data items | 10a | List and define all outcomes for which data were sought. Specify whether all results that were compatible with each outcome domain in each study were sought (e.g. for all measures, time points, analyses), and if not, the methods used to decide which results to collect. | Method, Introduction |
| | 10b | List and define all other variables for which data were sought (e.g. participant and intervention characteristics, funding sources). Describe any assumptions made about any missing or unclear information. | Method |
| Study risk of bias assessment | 11 | Specify the methods used to assess risk of bias in the included studies, including details of the tool(s) used, and if applicable, details of automation tools used in the process. | Method, *SM* 3 |
| Effect measures | 12 | Specify for each outcome the effect measure(s) (e.g. risk ratio, mean difference) used in the synthesis or presentation of results. | Method |
| Synthesis methods | 13a | Describe the processes used to decide which studies were eligible for each synthesis (e.g. tabulating the study intervention characteristics and comparing against the planned groups for each synthesis (item #5)). | Method |
| | 13b | Describe any methods required to prepare the data for presentation or synthesis, such as handling of missing summary statistics, or data conversions. | Method |
| | 13c | Describe any methods used to tabulate or visually display results of individual studies and syntheses. | Method |
| | 13d | Describe any methods used to synthesize results and provide a rationale for the choice(s). If meta-analysis was performed, describe the model(s), method(s) to identify the presence and extent of statistical heterogeneity, and software package(s) used. | Method |
| | 13e | Describe any methods used to explore possible causes of heterogeneity among study results (e.g. subgroup analysis, meta-regression). | Method |
| | 13f | Describe any sensitivity analyses conducted to assess robustness of the synthesized results. | Method |
| Reporting bias assessment | 14 | Describe any methods used to assess risk of bias due to missing results in a synthesis (arising from reporting biases). | Method |
| Certainty assessment | 15 | Describe any methods used to assess certainty (or confidence) in the body of evidence for an outcome. | Method |

| Section and Topic | Item # | Checklist item | Location where item is reported |
|---|---|---|---|
| **RESULTS** | | | |
| Study selection | 16a | Describe the results of the search and selection process, from the number of records identified in the search to the number of studies included in the review, ideally using a flow diagram. | Method, Results |
| | 16b | Cite studies that might appear to meet the inclusion criteria, but which were excluded, and explain why they were excluded. | Method, Discussion |
| Study characteristics | 17 | Cite each included study and present its characteristics. | Results |
| Risk of bias in studies | 18 | Present assessments of risk of bias for each included study. | Results |
| Results of individual studies | 19 | For all outcomes, present, for each study: (a) summary statistics for each group (where appropriate) and (b) an effect estimate and its precision (e.g. confidence/credible interval), ideally using structured tables or plots. | Results |
| Results of syntheses | 20a | For each synthesis, briefly summarise the characteristics and risk of bias among contributing studies. | Results |
| | 20b | Present results of all statistical syntheses conducted. If meta-analysis was done, present for each the summary estimate and its precision (e.g. confidence/credible interval) and measures of statistical heterogeneity. If comparing groups, describe the direction of the effect. | Results |
| | 20c | Present results of all investigations of possible causes of heterogeneity among study results. | Results |
| | 20d | Present results of all sensitivity analyses conducted to assess the robustness of the synthesized results. | Results |
| Reporting biases | 21 | Present assessments of risk of bias due to missing results (arising from reporting biases) for each synthesis assessed. | Results |
| Certainty of evidence | 22 | Present assessments of certainty (or confidence) in the body of evidence for each outcome assessed. | Results |
| **DISCUSSION** | | | |
| Discussion | 23a | Provide a general interpretation of the results in the context of other evidence. | Discussion |
| | 23b | Discuss any limitations of the evidence included in the review. | Discussion |
| | 23c | Discuss any limitations of the review processes used. | Discussion |
| | 23d | Discuss implications of the results for practice, policy, and future research. | Discussion |
| **OTHER INFORMATION** | | | |
| Registration and protocol | 24a | Provide registration information for the review, including register name and registration number, or state that the review was not registered. | NA |
| | 24b | Indicate where the review protocol can be accessed, or state that a protocol was not prepared. | Method |
| | 24c | Describe and explain any amendments to information provided at registration or in the protocol. | *SM* 12 |
| Support | 25 | Describe sources of financial or non-financial support for the review, and the role of the funders or sponsors in the review. | Acknowledge ment |
| Competing interests | 26 | Declare any competing interests of review authors. | Declaration of competing interest |
| Availability of data, code and other materials | 27 | Report which of the following are publicly available and where they can be found: template data collection forms; data extracted from included studies; data used for all analyses; analytic code; any other materials used in the review. | Data and code availability |

#### 2 Study characteristics and effect size

Study characteristics and effect size included were reported in separate csv files. All datasets were permanently saved in the OSF repository: https://osf.io/9mh5d/. Replication is allowed.

Table S2.1 contains all demographic information, publication details, as well as summarized moderators and results. Tables S2.2-2.5 are sub-tables for coupling phase, SP amplitude, coupling strength, and coupling percentage, respectively. Each contains detailed physiological and behavioral conditions for each effect size, mean values of each sleep measure, and effect sizes.

#### 3 Risk of Bias Assessment

Table S 3.1: Risk of Bias Assessment Supplemental Criteria

| **Domain** | **Supplemental Signaling Questions** |
|---|---|
| Bias due to confounding variables | 1.1 Was sufficient information provided to assess the presence of major potential confounding variables?<br>1.2 Were major potential confounding variables not relevant to study controlled during the data collection and analysis? Were influences from other experimental tasks or stimuli existed?<br>1.3 Were confounding factors (such as gender, age, etc.) added to models to calculate and interpret as the "corrected" effect size? |
| Bias due to subject selection | 2.1 Was subject selection representative? Can subjects represent the population or community targeted by the experiment?<br>2.2 Was a random sampling method applied during data collection? Are subjects recruited mostly from a single source (e.g. university) or at different times and caused biases?<br>2.3 Are the subjects independent from each other? Are the subjects socially connected (e.g. patients and their relatives, between groups if there are multiple groups in the original study)? Have the same subjects been measured repeatedly in pretest-posttest designs? |
| Bias due to classification of groups | 3.1 For studies with multiple groups, can group(s) containing only healthy human subjects without intervention be clearly classified?<br>3.2 When reporting the effect size, did the authors report the effect size separately for different groups? |
| Bias due to missing outcome data | 4.1 Were only scatterplots reported in the article, necessitating the use of graph tools to estimate the effect size?<br>4.2 Was data including $t$ statistics, $p$-values, $\beta$ statistics, and $\eta^2$ the only data provided that could be used to estimate the correlation, or only imprecise data provided for non-significant correlation?<br>4.3 Have pre-sleep/sleep/post-sleep memories and/or sleep data for individual participants been lost? If true, were missing outcome data interpolated, averaged, simulated, or deleted? |

| Domain | Supplemental Signaling Questions |
| --- | --- |
| Bias in measurement of the outcome | 5.1 Were measurements, units and signal processing approaches (including slow-oscillation, spindle, and coupling detection), used by the paper reliable and consistent with others?<br>5.2 Were only nonparametric effect sizes including spearman's rho ($\rho$), kendall's tau ($\tau$) or subjective estimation reported instead of Pearson's $r$, Fisher's z, or circular-linear $r$?<br>5.3 Did the authors analyze, transform, or clarify non-normal data, introduce resampling techniques and/or exclude outliers? |
| Bias in selection of the reported result | 6.1 For multiple groups measured, if only group(s) with the largest effect size, or supported their hypotheses were reported, while non-significant or contradictory results were omitted?<br>6.2 Have the authors declared their research proposal and expected outcomes within the framework of pre-registration, or declared any conflict of interest in the paper? |
| Overall Bias | 7.1 Does the article not fully meet the applicable expectations of the Risk Of Bias In Non-randomized Studies of Interventions (Robins-I) and the above supplementary criteria in multiple domains listed?<br>7.2 Alternatively, does the article significantly conflict with the criteria in one of these domains? |

Due to the absence of a dedicated risk of bias assessment tool or guideline specifically designed for meta-analysis of correlation coefficients, we made minor adaptations based on established criteria and signaling questions from the Risk Of Bias In Non-randomized Studies of Interventions (Robins-I) framework[1,2]. These adaptations entailed substituting descriptions related to interventions and control experiments with references to sleep and single-group experiments, omitting criteria that were not applicable to the specific objectives of our study, adjusting domain classifications, and included standards from the NIH Study Quality Assessment Tools for Before-After (Pre-Post) Studies With No Control Group[3] in the evaluation of certain studies. In addition, we have augmented our assessment comprehensively by introducing supplemental signaling questions, as outlined in Table S3.2, in accordance with the research methods and data extraction procedures detailed in the meta-analysis. For the data set we requested from the author, it would still be marked as low risk when the above conditions were met. The full version of signaling questions can be found in Robins-I. The outcomes of the risk of bias assessment for each research paper included in the meta-analysis can be found in Figure S3.1, generated by the "robvis" package[4] in the R statistical computing environment[5].

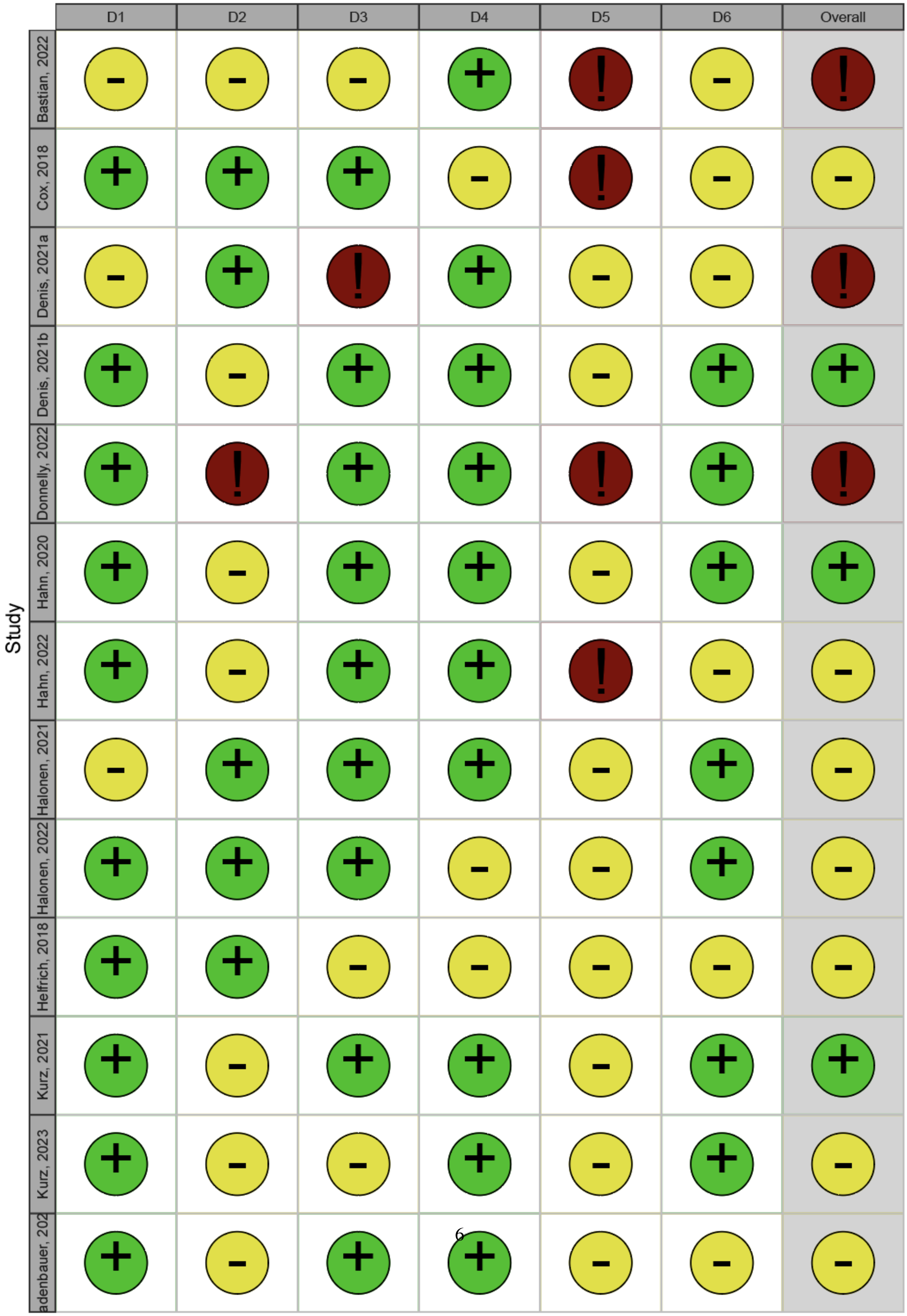
D1
D2
D3
D4
D5
D6
Overall
Study
Bastian, 2022
Cox, 2018
Denis, 2021a
Denis, 2021b
Donnelly, 2022
Hahn, 2020
Hahn, 2022
Halonen, 2021
Halonen, 2022
Helfrich, 2018
Kurz, 2021
Kurz, 2023
adenbauer, 202

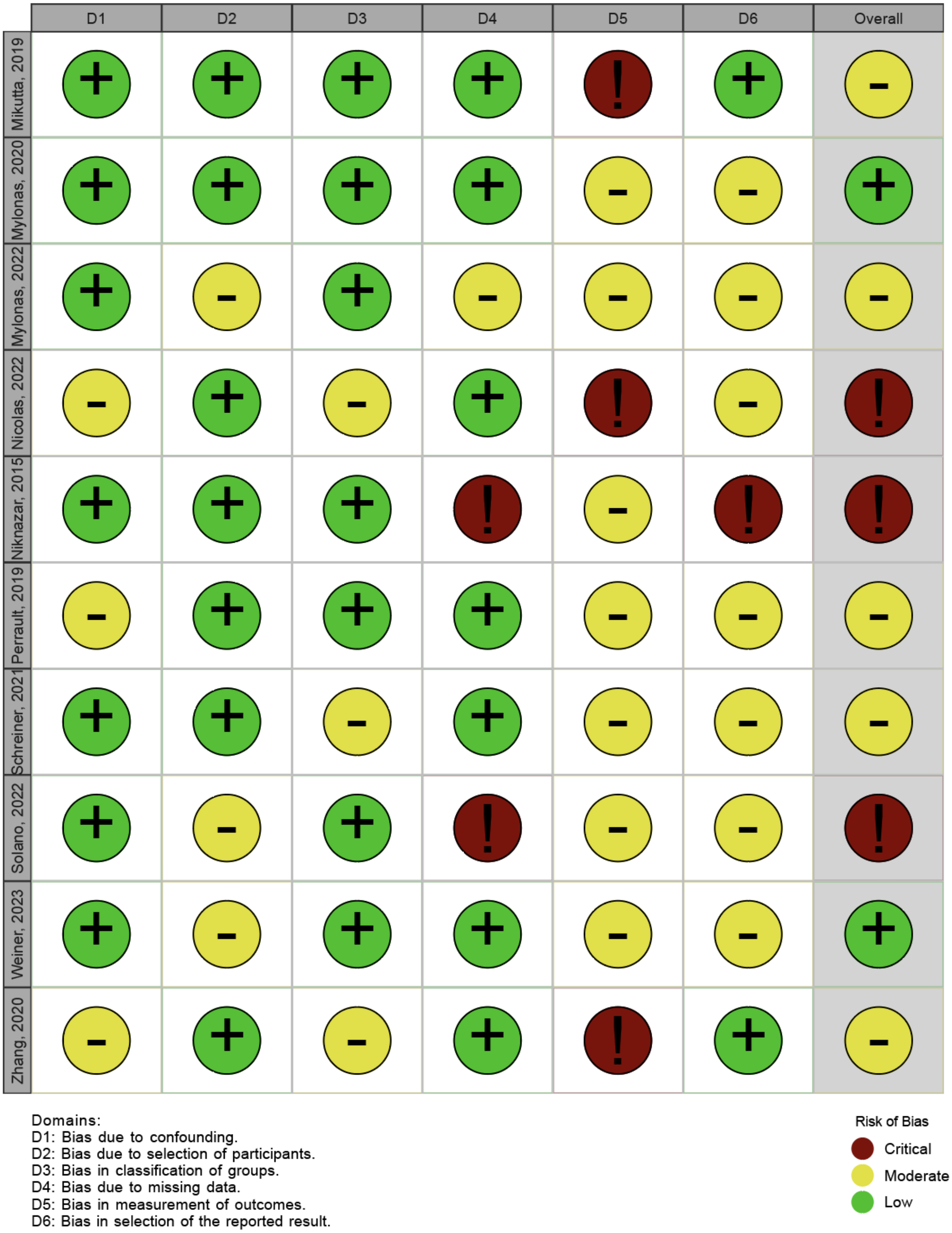


Figure S 3.1: Risk of bias (ROB) assessment for individual studies

#### 4 Model Diagnostics

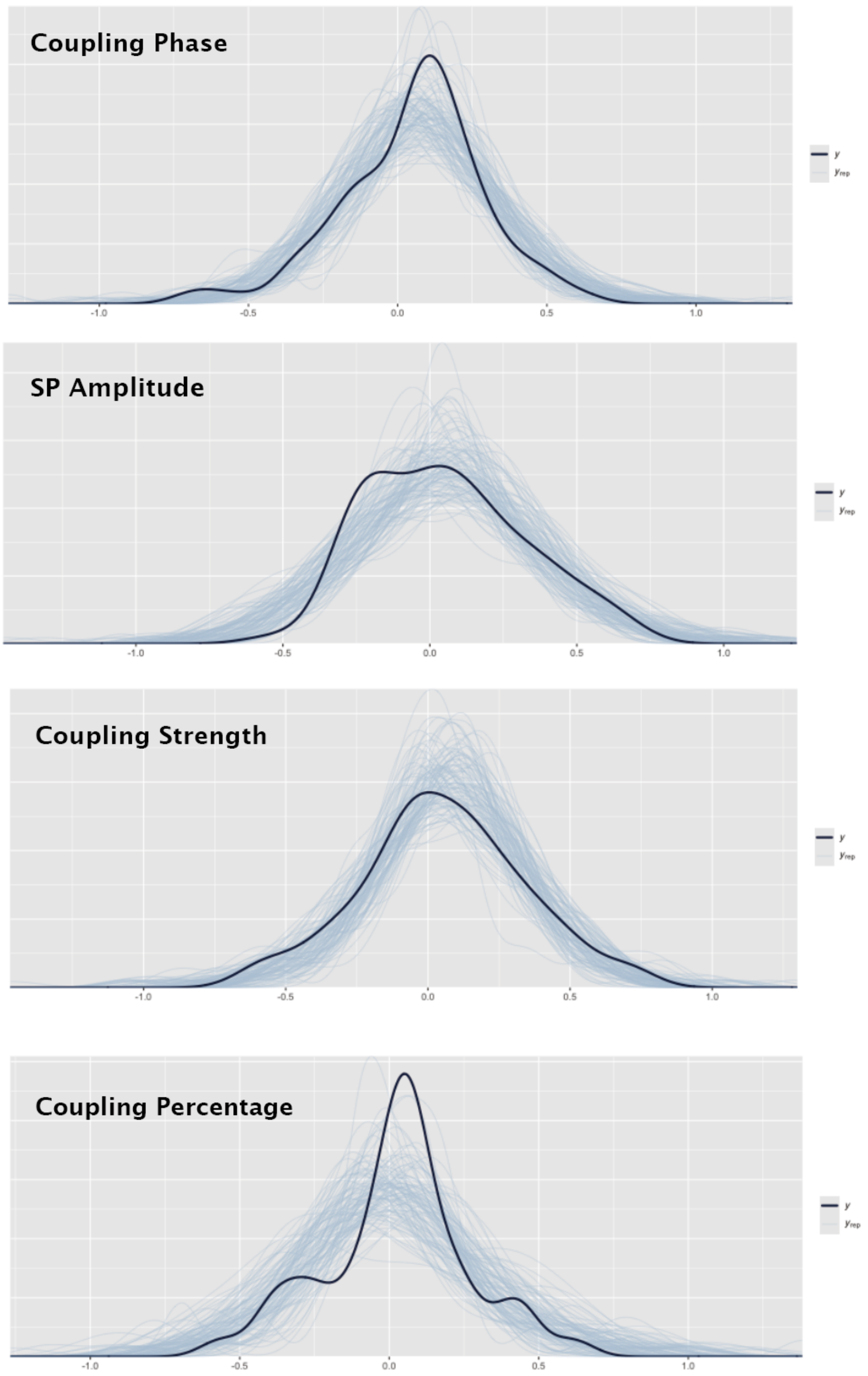




Figure S 4.1: Posterior predictive check of overall models of coupling-memory association measures

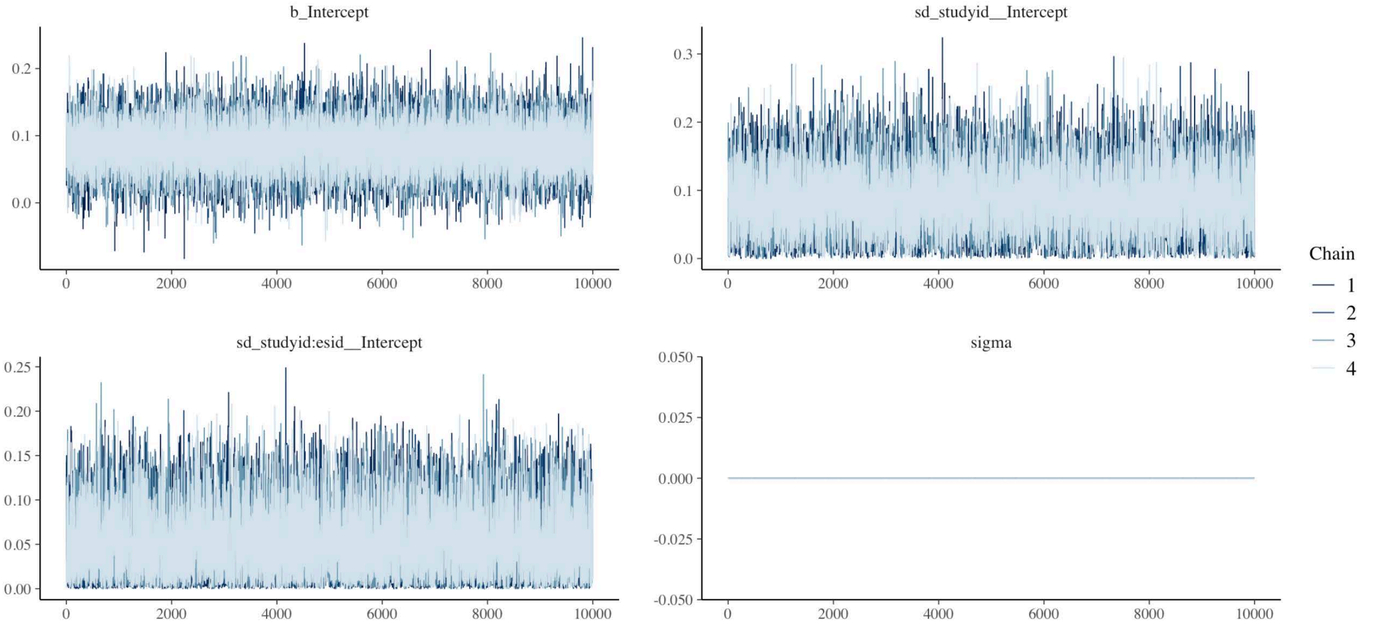


Figure S 4.2: Example of the trace plot used to assess the convergence of models. Well-mixed chains showing no trends or drifts over iterations indicate successful convergence.

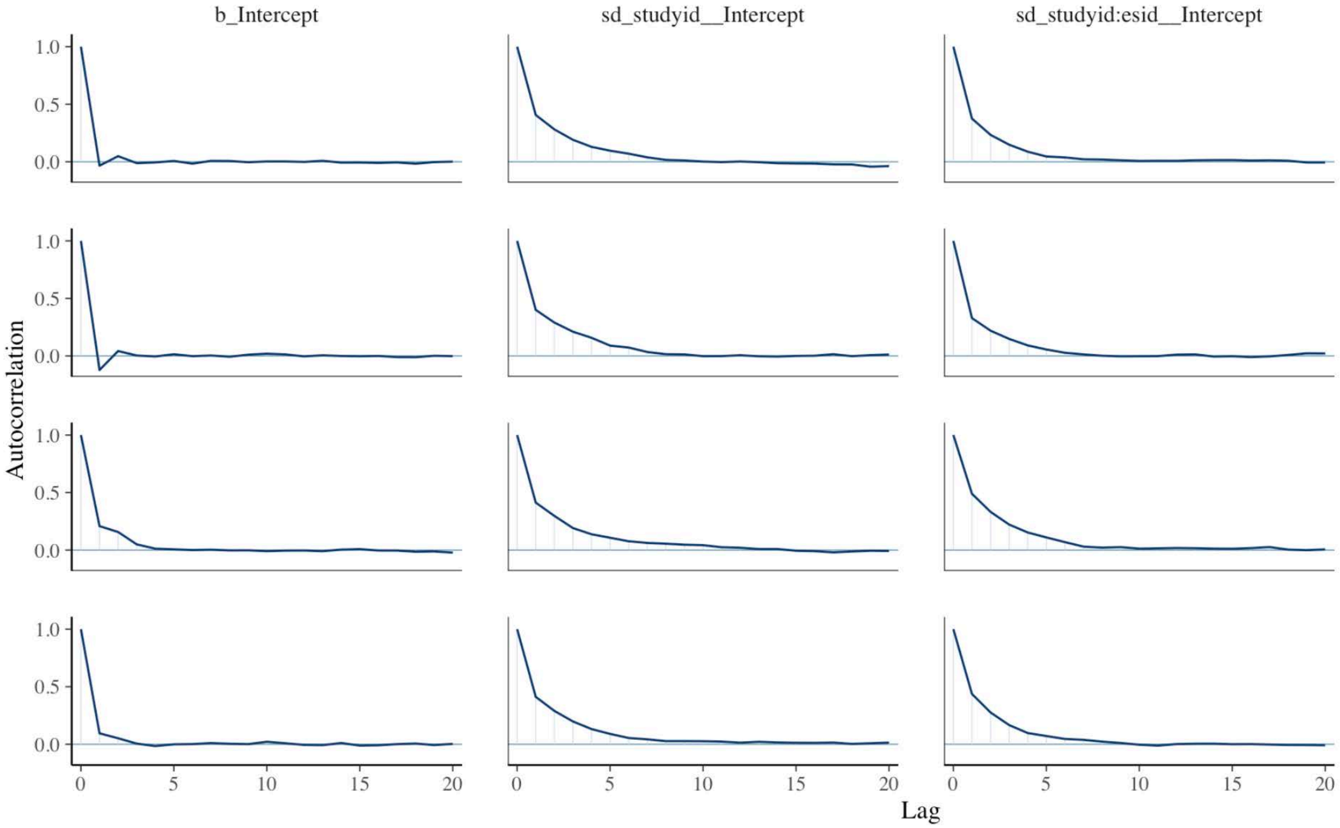


Figure S 4.3: Example of the autocorrelation plot used to assess the independence of samples across iterations, with a rapid decay of autocorrelation indicating efficient sampling and low within-chain redundancy.

### 5 Effect size-level forest plot

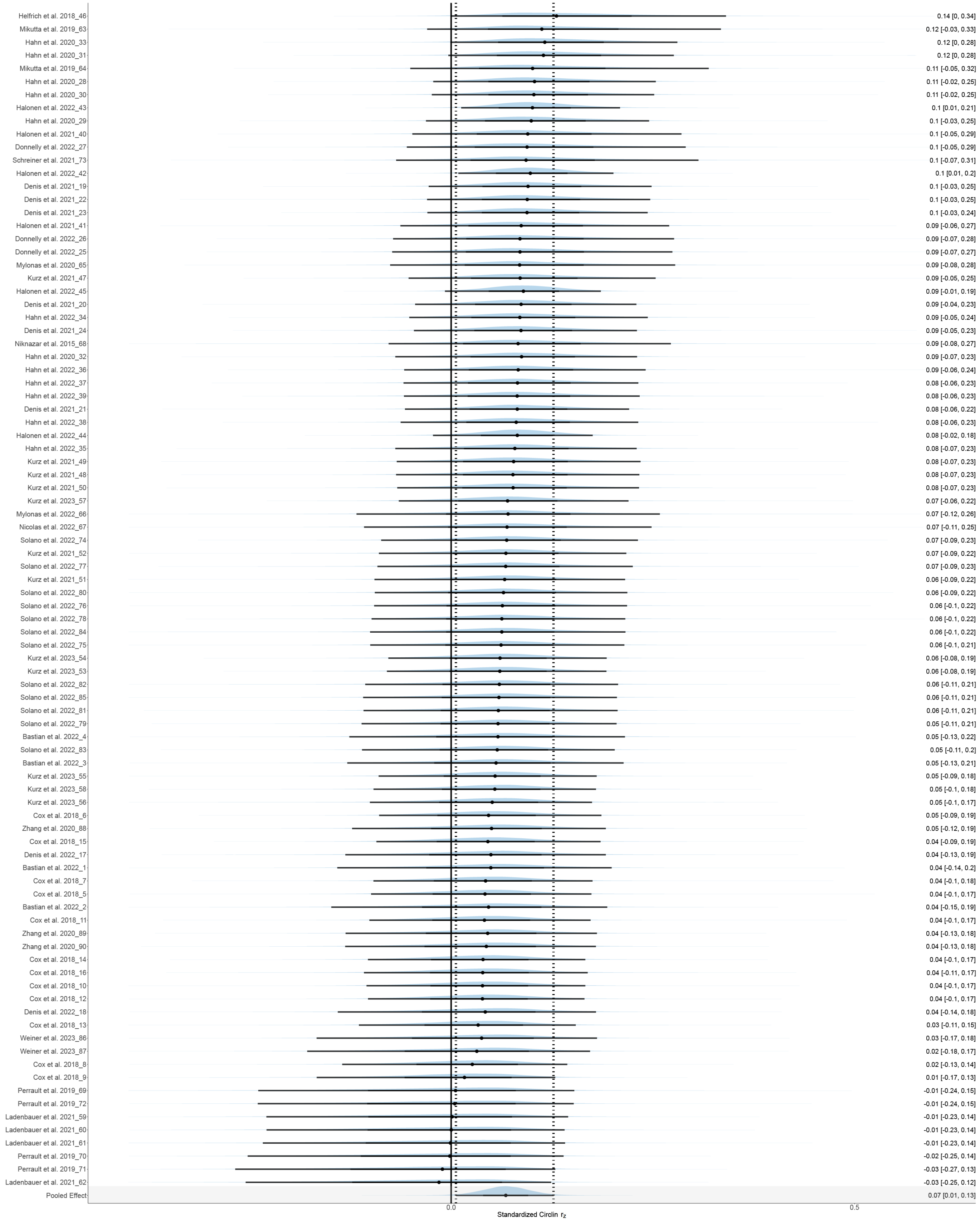


Figure S 5.1: Effect size-level forest plot for the overall model of coupling phase

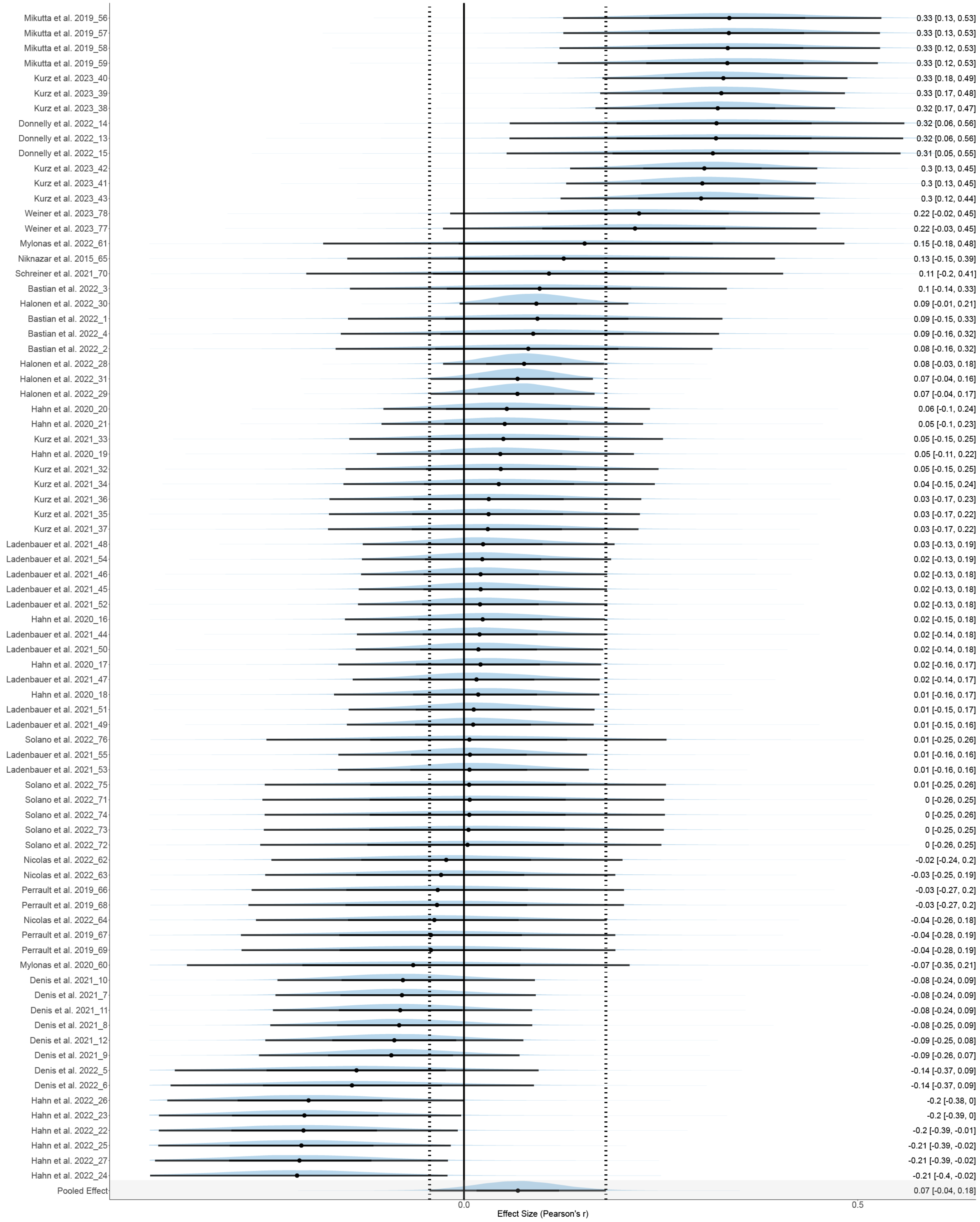


Figure S 5.2: Effect size-level forest plot for the overall model of spindle amplitude

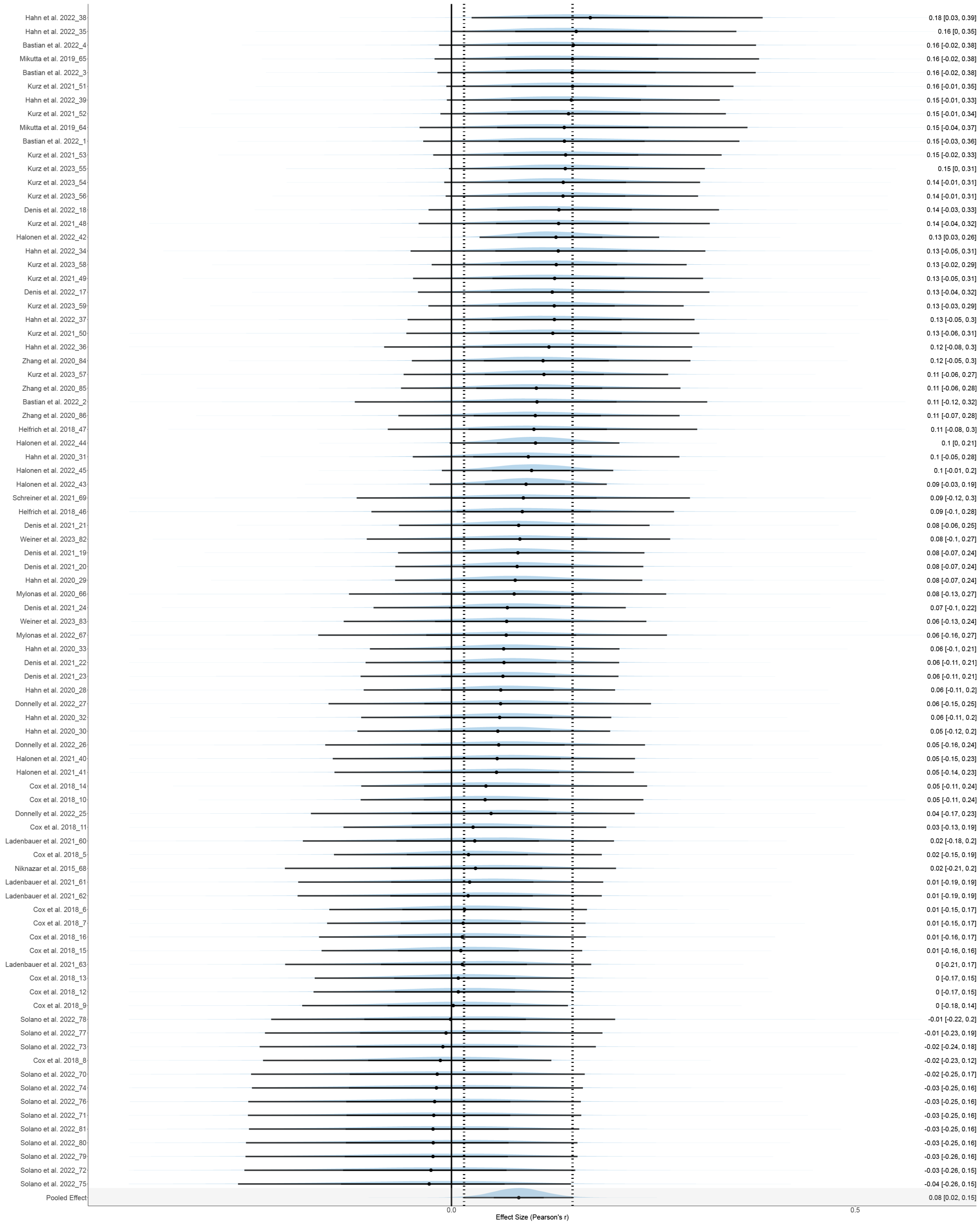

Figure S 5.3: Effect size-level forest plot for the overall model of coupling strength

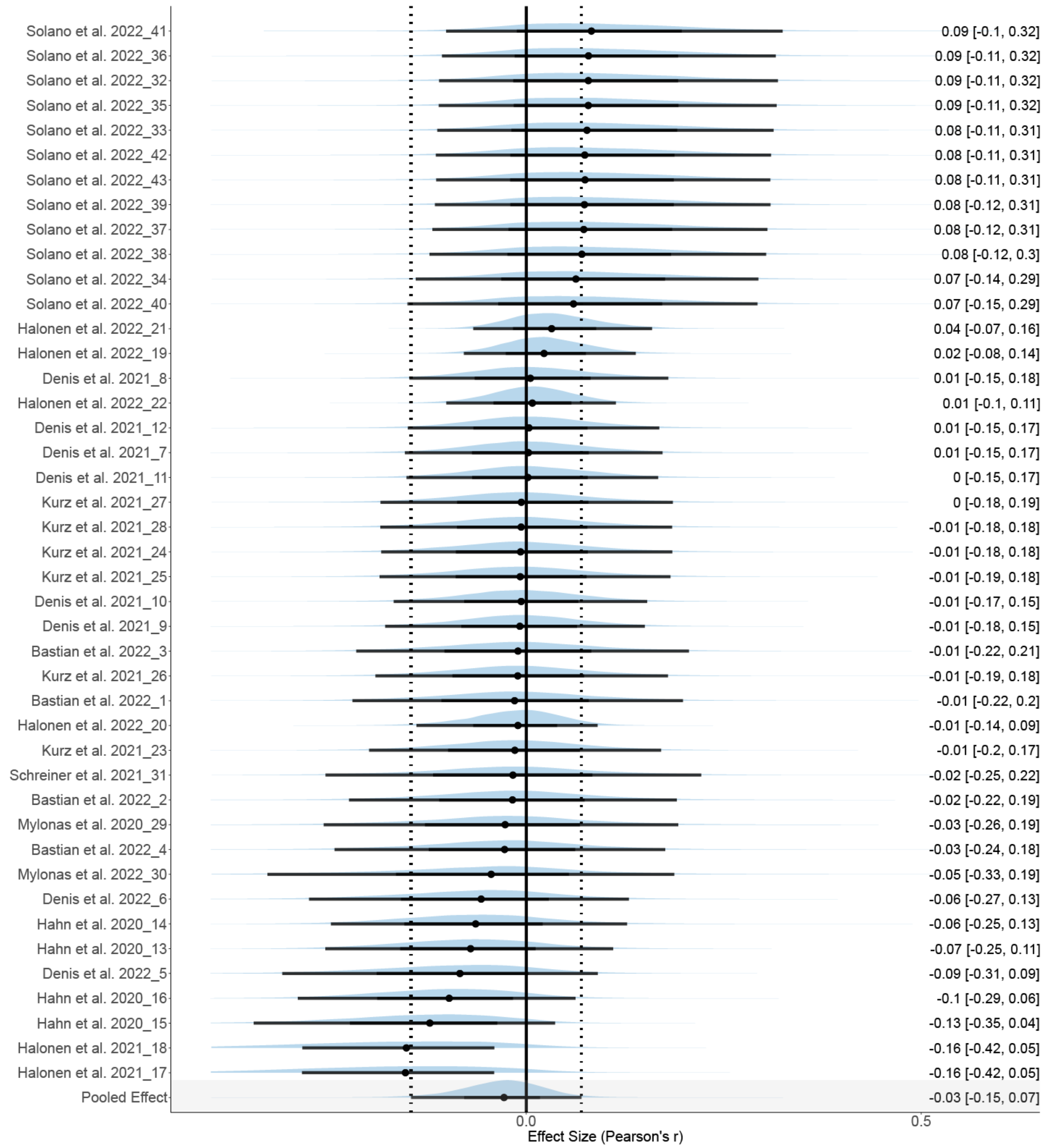


Figure S 5.4: Effect size-level forest plot for the overall model of coupling percentage

Notes. Overall model forest plot at the effect size-level. The solid vertical line represents the mean correlation coefficient under the null hypothesis. Dashed lines indicate the 95% credible interval ($CrI$) of the pooled effect size. The black point and error bar for each effect size shows the adjusted estimation of effect size and 95% $CrI$ combining data and prior information, with corresponding values reported in the right side of the figure. On the left, numbers reported after publication years correspond to the esid column labeled in the dataset S2.2-2.5.

### 6 Pareto *K* diagnosis statistics

| Pareto k diagnosis statistics for the phase model | | | | | | | | |
|---|---|---|---|---|---|---|---|---|
| ESID | Study | Pareto k | ESID | Study | Pareto k | ESID | Study | Pareto k |
| 1 | Bastian et al. 2022 | 0.204 | 31 | Hahn et al. 2020 | 0.401 | 61 | Ladenbauer et al. 2021 | 0.324 |
| 2 | Bastian et al. 2022 | 0.221 | 32 | Hahn et al. 2020 | 0.455 | 62 | Ladenbauer et al. 2021 | 0.358 |
| 3 | Bastian et al. 2022 | 0.282 | 33 | Hahn et al. 2020 | 0.571 | 63 | Mikutta et al. 2019 | 0.397 |
| 4 | Bastian et al. 2022 | 0.247 | 34 | Hahn et al. 2022 | 0.273 | 64 | Mikutta et al. 2019 | 0.229 |
| 5 | Cox et al. 2018 | 0.339 | 35 | Hahn et al. 2022 | 0.185 | 65 | Mylonas et al. 2020 | 0.399 |
| 6 | Cox et al. 2018 | 0.401 | 36 | Hahn et al. 2022 | 0.376 | 66 | Mylonas et al. 2022 | 0.395 |
| 7 | Cox et al. 2018 | 0.304 | 37 | Hahn et al. 2022 | 0.332 | 67 | Nicolas et al. 2022 | 0.434 |
| 8 | Cox et al. 2018 | 0.259 | 38 | Hahn et al. 2022 | 0.367 | 68 | Niknazar et al. 2015 | 0.489 |
| 9 | Cox et al. 2018 | 0.255 | 39 | Hahn et al. 2022 | 0.222 | 69 | Perrault et al. 2019 | 0.320 |
| 10 | Cox et al. 2018 | 0.296 | 40 | Halonen et al. 2021 | 0.453 | 70 | Perrault et al. 2019 | 0.237 |
| 11 | Cox et al. 2018 | 0.357 | 41 | Halonen et al. 2021 | 0.417 | 71 | Perrault et al. 2019 | 0.345 |
| 12 | Cox et al. 2018 | 0.279 | 42 | Halonen et al. 2022 | 0.705 * | 72 | Perrault et al. 2019 | 0.210 |
| 13 | Cox et al. 2018 | 0.377 | 43 | Halonen et al. 2022 | 0.498 | 73 | Schreiner et al. 2021 | 0.391 |
| 14 | Cox et al. 2018 | 0.421 | 44 | Halonen et al. 2022 | 0.52 | 74 | Solano et al. 2022 | 0.103 |
| 15 | Cox et al. 2018 | 0.356 | 45 | Halonen et al. 2022 | 0.551 | 75 | Solano et al. 2022 | 0.166 |
| 16 | Cox et al. 2018 | 0.251 | 46 | Helfrich et al. 2018 | 0.531 | 76 | Solano et al. 2022 | 0.195 |
| 17 | Denis et al. 2022 | 0.432 | 47 | Kurz et al. 2021 | 0.365 | 77 | Solano et al. 2022 | 0.125 |
| 18 | Denis et al. 2022 | 0.403 | 48 | Kurz et al. 2021 | 0.255 | 78 | Solano et al. 2022 | 0.130 |
| 19 | Denis et al. 2021 | 0.331 | 49 | Kurz et al. 2021 | 0.328 | 79 | Solano et al. 2022 | 0.102 |
| 20 | Denis et al. 2021 | 0.324 | 50 | Kurz et al. 2021 | 0.322 | 80 | Solano et al. 2022 | 0.187 |
| 21 | Denis et al. 2021 | 0.373 | 51 | Kurz et al. 2021 | 0.285 | 81 | Solano et al. 2022 | 0.187 |
| 22 | Denis et al. 2021 | 0.380 | 52 | Kurz et al. 2021 | 0.369 | 82 | Solano et al. 2022 | 0.141 |
| 23 | Denis et al. 2021 | 0.235 | 53 | Kurz et al. 2023 | 0.289 | 83 | Solano et al. 2022 | 0.167 |
| 24 | Denis et al. 2021 | 0.403 | 54 | Kurz et al. 2023 | 0.304 | 84 | Solano et al. 2022 | 0.291 |
| 25 | Donnelly et al. 2022 | 0.253 | 55 | Kurz et al. 2023 | 0.304 | 85 | Solano et al. 2022 | 0.197 |
| 26 | Donnelly et al. 2022 | 0.288 | 56 | Kurz et al. 2023 | 0.413 | 86 | Weiner et al. 2023 | 0.340 |
| 27 | Donnelly et al. 2022 | 0.278 | 57 | Kurz et al. 2023 | 0.324 | 87 | Weiner et al. 2023 | 0.347 |
| 28 | Hahn et al. 2020 | 0.305 | 58 | Kurz et al. 2023 | 0.224 | 88 | Zhang et al. 2020 | 0.281 |
| 29 | Hahn et al. 2020 | 0.322 | 59 | Ladenbauer et al. 2021 | 0.272 | 89 | Zhang et al. 2020 | 0.342 |
| 30 | Hahn et al. 2020 | 0.293 | 60 | Ladenbauer et al. 2021 | 0.288 | 90 | Zhang et al. 2020 | 0.395 |

Figure S 6.1: Pareto *k* diagnostic statistics for the coupling phase model

| Pareto k diagnosis statistics for the SP amplitude model | | | | | | | | |
|---|---|---|---|---|---|---|---|---|
| ESID | Study | Pareto k | ESID | Study | Pareto k | ESID | Study | Pareto k |
| 1 | Bastian et al. 2022 | 0.224 | 27 | Hahn et al. 2022 | 0.303 | 53 | Ladenbauer et al. 2021 | 0.357 |
| 2 | Bastian et al. 2022 | 0.173 | 28 | Halonen et al. 2022 | 0.658 | 54 | Ladenbauer et al. 2021 | 0.283 |
| 3 | Bastian et al. 2022 | 0.291 | 29 | Halonen et al. 2022 | 0.591 | 55 | Ladenbauer et al. 2021 | 0.407 |
| 4 | Bastian et al. 2022 | 0.195 | 30 | Halonen et al. 2022 | 0.529 | 56 | Mikutta et al. 2019 | 0.292 |
| 5 | Denis et al. 2022 | 0.453 | 31 | Halonen et al. 2022 | 0.737 * | 57 | Mikutta et al. 2019 | 0.313 |
| 6 | Denis et al. 2022 | 0.489 | 32 | Kurz et al. 2021 | 0.354 | 58 | Mikutta et al. 2019 | 0.321 |
| 7 | Denis et al. 2021 | 0.411 | 33 | Kurz et al. 2021 | 0.302 | 59 | Mikutta et al. 2019 | 0.270 |
| 8 | Denis et al. 2021 | 0.389 | 34 | Kurz et al. 2021 | 0.247 | 60 | Mylonas et al. 2020 | 0.581 |
| 9 | Denis et al. 2021 | 0.324 | 35 | Kurz et al. 2021 | 0.345 | 61 | Mylonas et al. 2022 | 0.576 |
| 10 | Denis et al. 2021 | 0.342 | 36 | Kurz et al. 2021 | 0.342 | 62 | Nicolas et al. 2022 | 0.401 |
| 11 | Denis et al. 2021 | 0.427 | 37 | Kurz et al. 2021 | 0.252 | 63 | Nicolas et al. 2022 | 0.347 |
| 12 | Denis et al. 2021 | 0.371 | 38 | Kurz et al. 2023 | 0.311 | 64 | Nicolas et al. 2022 | 0.393 |
| 13 | Donnelly et al. 2022 | 0.413 | 39 | Kurz et al. 2023 | 0.286 | 65 | Niknazar et al. 2015 | 0.568 |
| 14 | Donnelly et al. 2022 | 0.381 | 40 | Kurz et al. 2023 | 0.426 | 66 | Perrault et al. 2019 | 0.309 |
| 15 | Donnelly et al. 2022 | 0.336 | 41 | Kurz et al. 2023 | 0.485 | 67 | Perrault et al. 2019 | 0.218 |
| 16 | Hahn et al. 2020 | 0.500 | 42 | Kurz et al. 2023 | 0.339 | 68 | Perrault et al. 2019 | 0.265 |
| 17 | Hahn et al. 2020 | 0.403 | 43 | Kurz et al. 2023 | 0.353 | 69 | Perrault et al. 2019 | 0.300 |
| 18 | Hahn et al. 2020 | 0.428 | 44 | Ladenbauer et al. 2021 | 0.298 | 70 | Schreiner et al. 2021 | 0.542 |
| 19 | Hahn et al. 2020 | 0.442 | 45 | Ladenbauer et al. 2021 | 0.352 | 71 | Solano et al. 2022 | 0.120 |
| 20 | Hahn et al. 2020 | 0.458 | 46 | Ladenbauer et al. 2021 | 0.334 | 72 | Solano et al. 2022 | 0.288 |
| 21 | Hahn et al. 2020 | 0.442 | 47 | Ladenbauer et al. 2021 | 0.281 | 73 | Solano et al. 2022 | 0.167 |
| 22 | Hahn et al. 2022 | 0.288 | 48 | Ladenbauer et al. 2021 | 0.203 | 74 | Solano et al. 2022 | 0.151 |
| 23 | Hahn et al. 2022 | 0.277 | 49 | Ladenbauer et al. 2021 | 0.469 | 75 | Solano et al. 2022 | 0.180 |
| 24 | Hahn et al. 2022 | 0.391 | 50 | Ladenbauer et al. 2021 | 0.388 | 76 | Solano et al. 2022 | 0.165 |
| 25 | Hahn et al. 2022 | 0.286 | 51 | Ladenbauer et al. 2021 | 0.229 | 77 | Weiner et al. 2023 | 0.358 |
| 26 | Hahn et al. 2022 | 0.378 | 52 | Ladenbauer et al. 2021 | 0.3 | 78 | Weiner et al. 2023 | 0.389 |

Figure S 6.2: Pareto *k* diagnostic statistics for the spindle amplitude model

| Pareto k diagnosis statistics for the strength model | | | | | | | | |
|---|---|---|---|---|---|---|---|---|
| ESID | Study | Pareto k | ESID | Study | Pareto k | ESID | Study | Pareto k |
| 1 | Bastian et al. 2022 | 0.315 | 30 | Hahn et al. 2020 | 0.485 | 59 | Kurz et al. 2023 | 0.313 |
| 2 | Bastian et al. 2022 | 0.434 | 31 | Hahn et al. 2020 | 0.425 | 60 | Ladenbauer et al. 2021 | 0.284 |
| 3 | Bastian et al. 2022 | 0.401 | 32 | Hahn et al. 2020 | 0.48 | 61 | Ladenbauer et al. 2021 | 0.260 |
| 4 | Bastian et al. 2022 | 0.404 | 33 | Hahn et al. 2020 | 0.464 | 62 | Ladenbauer et al. 2021 | 0.278 |
| 5 | Cox et al. 2018 | 0.345 | 34 | Hahn et al. 2022 | 0.271 | 63 | Ladenbauer et al. 2021 | 0.278 |
| 6 | Cox et al. 2018 | 0.459 | 35 | Hahn et al. 2022 | 0.348 | 64 | Mikutta et al. 2019 | 0.321 |
| 7 | Cox et al. 2018 | 0.394 | 36 | Hahn et al. 2022 | 0.281 | 65 | Mikutta et al. 2019 | 0.305 |
| 8 | Cox et al. 2018 | 0.332 | 37 | Hahn et al. 2022 | 0.318 | 66 | Mylonas et al. 2020 | 0.403 |
| 9 | Cox et al. 2018 | 0.271 | 38 | Hahn et al. 2022 | 0.475 | 67 | Mylonas et al. 2022 | 0.319 |
| 10 | Cox et al. 2018 | 0.363 | 39 | Hahn et al. 2022 | 0.473 | 68 | Niknazar et al. 2015 | 0.410 |
| 11 | Cox et al. 2018 | 0.221 | 40 | Halonen et al. 2021 | 0.349 | 69 | Schreiner et al. 2021 | 0.394 |
| 12 | Cox et al. 2018 | 0.396 | 41 | Halonen et al. 2021 | 0.334 | 70 | Solano et al. 2022 | 0.267 |
| 13 | Cox et al. 2018 | 0.436 | 42 | Halonen et al. 2022 | 0.703 * | 71 | Solano et al. 2022 | 0.228 |
| 14 | Cox et al. 2018 | 0.425 | 43 | Halonen et al. 2022 | 0.567 | 72 | Solano et al. 2022 | 0.261 |
| 15 | Cox et al. 2018 | 0.319 | 44 | Halonen et al. 2022 | 0.659 | 73 | Solano et al. 2022 | 0.191 |
| 16 | Cox et al. 2018 | 0.253 | 45 | Halonen et al. 2022 | 0.594 | 74 | Solano et al. 2022 | 0.187 |
| 17 | Denis et al. 2022 | 0.324 | 46 | Helfrich et al. 2018 | 0.441 | 75 | Solano et al. 2022 | 0.300 |
| 18 | Denis et al. 2022 | 0.394 | 47 | Helfrich et al. 2018 | 0.363 | 76 | Solano et al. 2022 | 0.197 |
| 19 | Denis et al. 2021 | 0.392 | 48 | Kurz et al. 2021 | 0.317 | 77 | Solano et al. 2022 | 0.187 |
| 20 | Denis et al. 2021 | 0.441 | 49 | Kurz et al. 2021 | 0.345 | 78 | Solano et al. 2022 | 0.222 |
| 21 | Denis et al. 2021 | 0.376 | 50 | Kurz et al. 2021 | 0.288 | 79 | Solano et al. 2022 | 0.300 |
| 22 | Denis et al. 2021 | 0.324 | 51 | Kurz et al. 2021 | 0.427 | 80 | Solano et al. 2022 | 0.209 |
| 23 | Denis et al. 2021 | 0.432 | 52 | Kurz et al. 2021 | 0.42 | 81 | Solano et al. 2022 | 0.285 |
| 24 | Denis et al. 2021 | 0.407 | 53 | Kurz et al. 2021 | 0.276 | 82 | Weiner et al. 2023 | 0.362 |
| 25 | Donnelly et al. 2022 | 0.248 | 54 | Kurz et al. 2023 | 0.413 | 83 | Weiner et al. 2023 | 0.414 |
| 26 | Donnelly et al. 2022 | 0.353 | 55 | Kurz et al. 2023 | 0.464 | 84 | Zhang et al. 2020 | 0.404 |
| 27 | Donnelly et al. 2022 | 0.197 | 56 | Kurz et al. 2023 | 0.412 | 85 | Zhang et al. 2020 | 0.380 |
| 28 | Hahn et al. 2020 | 0.477 | 57 | Kurz et al. 2023 | 0.347 | 86 | Zhang et al. 2020 | 0.404 |
| 29 | Hahn et al. 2020 | 0.496 | 58 | Kurz et al. 2023 | 0.376 | NA | NA | NA |

Figure S 6.3: Pareto *k* diagnostic statistics for the coupling strength model

| Pareto k diagnosis statistics for the percentage model | | | | | | | | |
|---|---|---|---|---|---|---|---|---|
| ESID | Study | Pareto k | ESID | Study | Pareto k | ESID | Study | Pareto k |
| 1 | Bastian et al. 2022 | 0.228 | 16 | Hahn et al. 2020 | 0.471 | 31 | Schreiner et al. 2021 | 0.488 |
| 2 | Bastian et al. 2022 | 0.251 | 17 | Halonen et al. 2021 | 0.479 | 32 | Solano et al. 2022 | 0.304 |
| 3 | Bastian et al. 2022 | 0.289 | 18 | Halonen et al. 2021 | 0.437 | 33 | Solano et al. 2022 | 0.308 |
| 4 | Bastian et al. 2022 | 0.385 | 19 | Halonen et al. 2022 | 0.814 * | 34 | Solano et al. 2022 | 0.159 |
| 5 | Denis et al. 2022 | 0.466 | 20 | Halonen et al. 2022 | 0.702 * | 35 | Solano et al. 2022 | 0.221 |
| 6 | Denis et al. 2022 | 0.524 | 21 | Halonen et al. 2022 | 0.543 | 36 | Solano et al. 2022 | 0.314 |
| 7 | Denis et al. 2021 | 0.406 | 22 | Halonen et al. 2022 | 0.754 * | 37 | Solano et al. 2022 | 0.255 |
| 8 | Denis et al. 2021 | 0.461 | 23 | Kurz et al. 2021 | 0.355 | 38 | Solano et al. 2022 | 0.284 |
| 9 | Denis et al. 2021 | 0.553 | 24 | Kurz et al. 2021 | 0.347 | 39 | Solano et al. 2022 | 0.332 |
| 10 | Denis et al. 2021 | 0.466 | 25 | Kurz et al. 2021 | 0.331 | 40 | Solano et al. 2022 | 0.237 |
| 11 | Denis et al. 2021 | 0.296 | 26 | Kurz et al. 2021 | 0.309 | 41 | Solano et al. 2022 | 0.167 |
| 12 | Denis et al. 2021 | 0.361 | 27 | Kurz et al. 2021 | 0.374 | 42 | Solano et al. 2022 | 0.319 |
| 13 | Hahn et al. 2020 | 0.295 | 28 | Kurz et al. 2021 | 0.322 | 43 | Solano et al. 2022 | 0.252 |
| 14 | Hahn et al. 2020 | 0.403 | 29 | Mylonas et al. 2020 | 0.462 | NA | NA | NA |
| 15 | Hahn et al. 2020 | 0.540 | 30 | Mylonas et al. 2022 | 0.493 | NA | NA | NA |

Figure S 6.4: Pareto *k* diagnostic statistics for the coupling percentage model

Notes. Pareto *k* can be interpreted in a similar way to Cook's distance in frequentist analysis. When an effect sizr has a Pareto *k* value larger than 0.7 or even 1.0, it can be classified as an influential data point and might introduce the issue of misspecification of the model. Consequently, we excluded all studies with Pareto *k* values larger than 0.7 from the focal analysis. ESID corresponds to the esid column labeled in the dataset S2.2-2.5.

### 7 Moderator models

Table S 7.1: Summary of models for SO-SP coupling-memory association measures

| Model | Moderator | Model Purposes |
|---|---|---|
| M | None | Study associations between SP amplitude, coupling phase, coupling strength, and coupling percentage, each in relation to memory consolidation. |
| M1 | Memory Task | Investigate potential distinctions in coupling and memory association mechanisms between declarative memory including verbal, spatial, emotional memory, as well as procedural memory. |
| M2 | Mean Age | Understand the potential impact of development and aging in coupling and memory associations. |
| M3 | Spindle Type | Explore how the coupling between SOs and fast or slow SPs predicts memory retention performance differently. |
| M4 | PSG Channel | Study the relation between sleep brain oscillations and memory in different cortical regions, as the frontal, central, and parietal areas were reported to be the most active area for SO-SP coupling but might play different roles. |
| M5 | Sleep Stage | Examine the impact of sleep stage on the relationship between SO-SP coupling and memory, considering that SPs are most active during N2 sleep, while SOs dominate cortical oscillation during SWS. |
| M6 | Sleep Phase | Investigate the potential impact of sleep timing and circadian rhythms on the relationship between SO-SP coupling and memory. |
| M7 | Age x Channel | Investigate interactions between age differences and PSG channels in the memory consolidation mechanism, considering the frontal lobe is the latest area of the brain to develop. |
| M8 | Age x Task | Study whether age increase implies a difference in predictive power of SO-SP coupling in the development of declarative and procedural memory consolidation. |
| M9 | Channel x Spindle | Examine interactions between SP types and cortical areas in models. |
| Mf | All predictors | Include all pre-specified moderators to the model as fixed effects to explore their explanatory power and potential collinearity. |
| Mc | None | Fitted controlled model for focal and sensitivity analysis. |

### 8 Posterior distribution of moderators

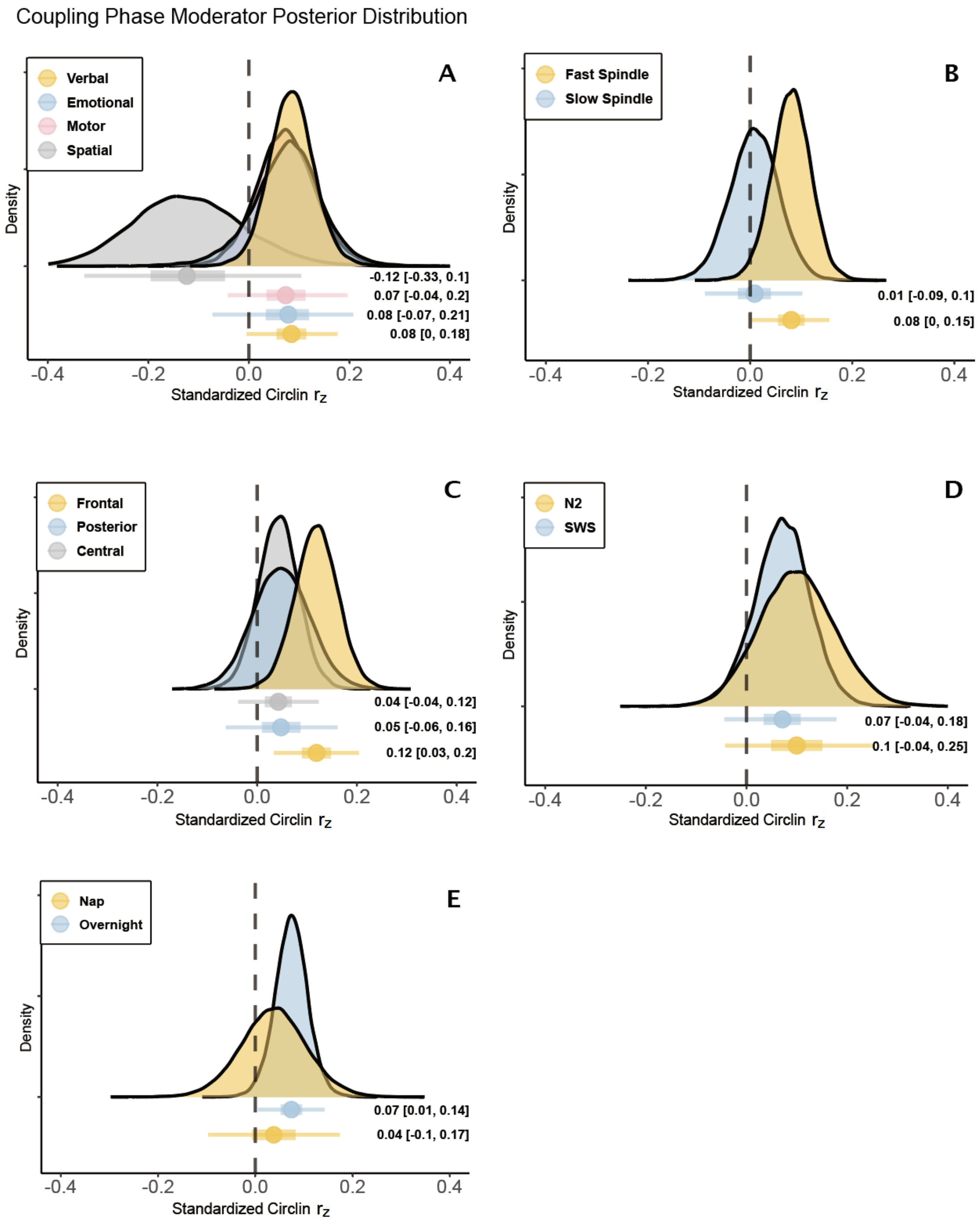




Figure S 8.1: Posterior distributions of moderation factors of the coupling phase

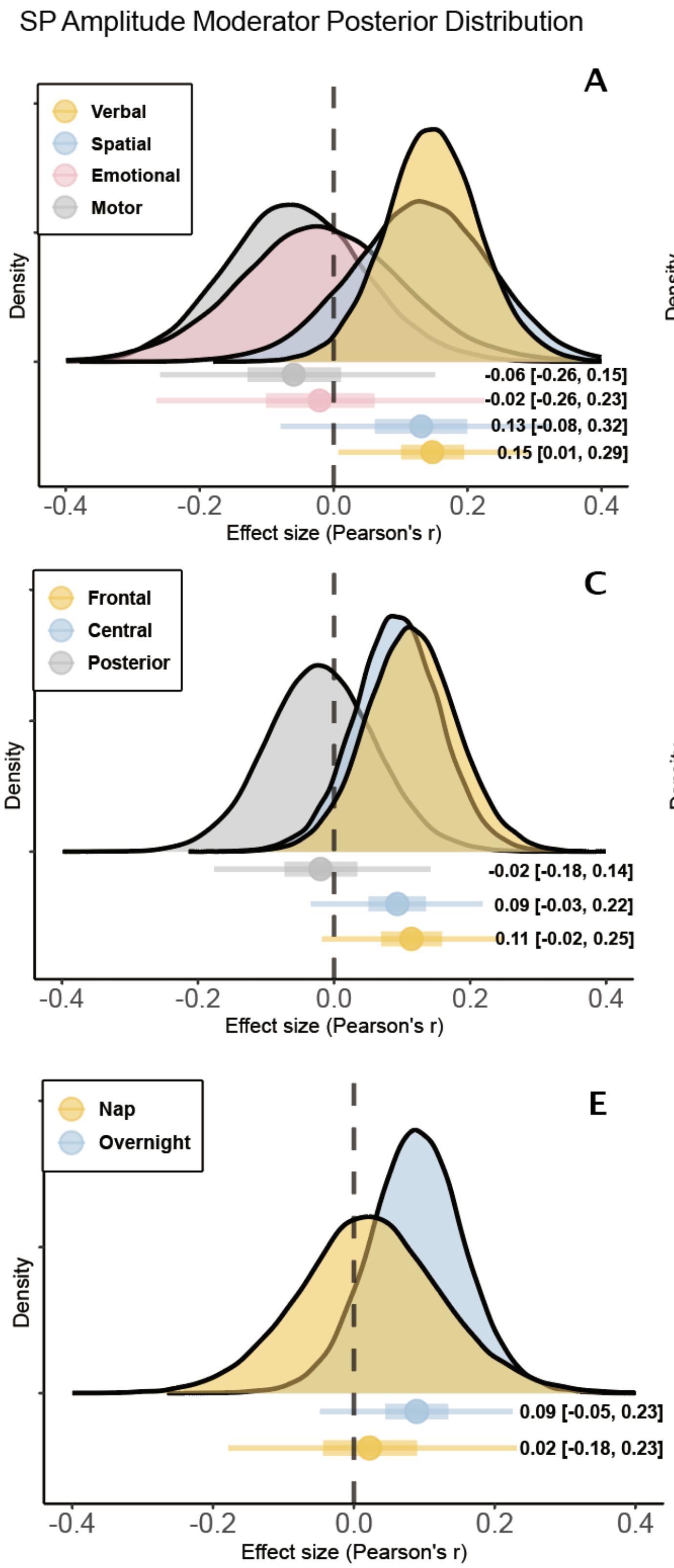


Figure S 8.2: Posterior distributions of moderation factors of the SP amplitude

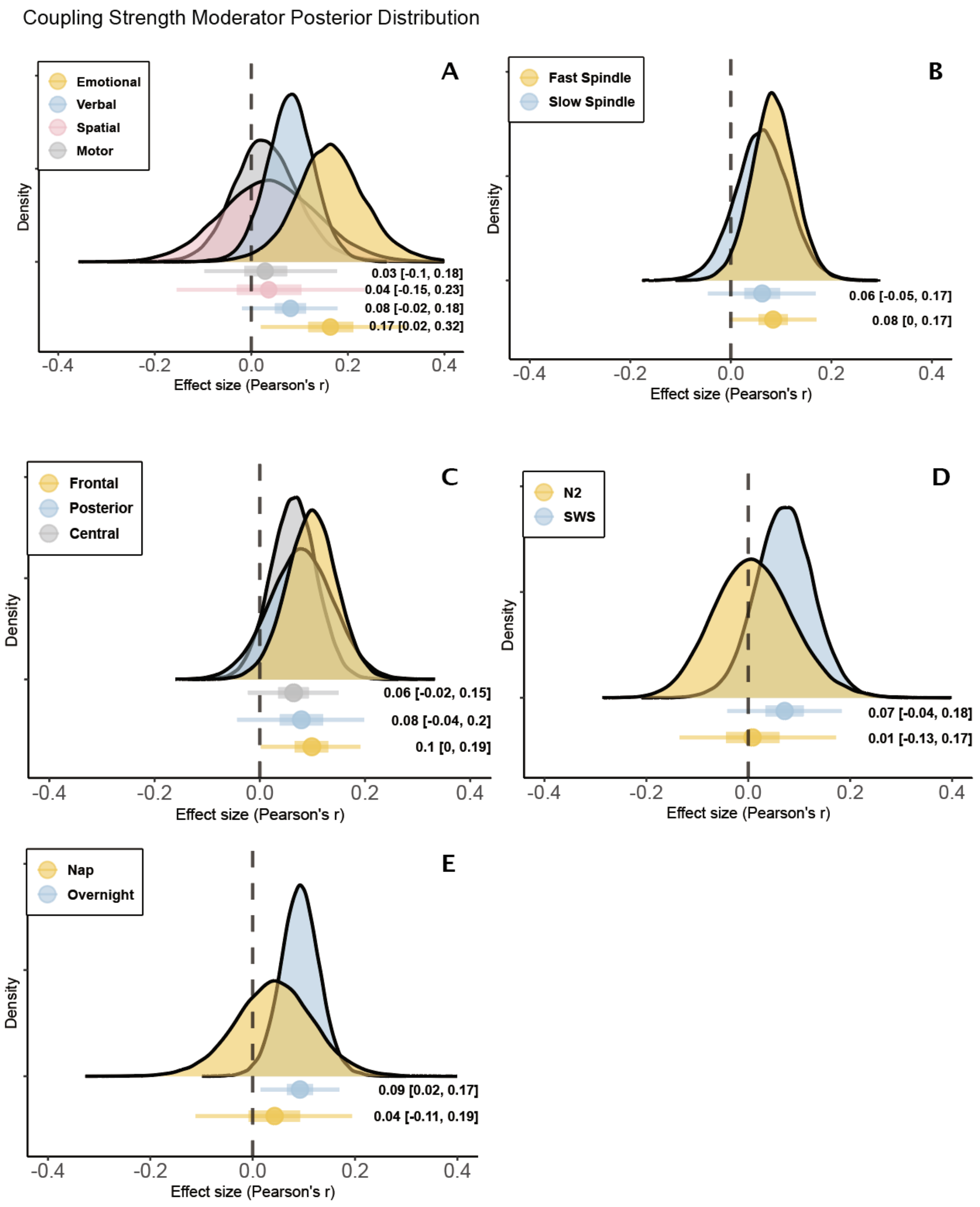


Figure S 8.3: Posterior distributions of moderation factors of the coupling strength

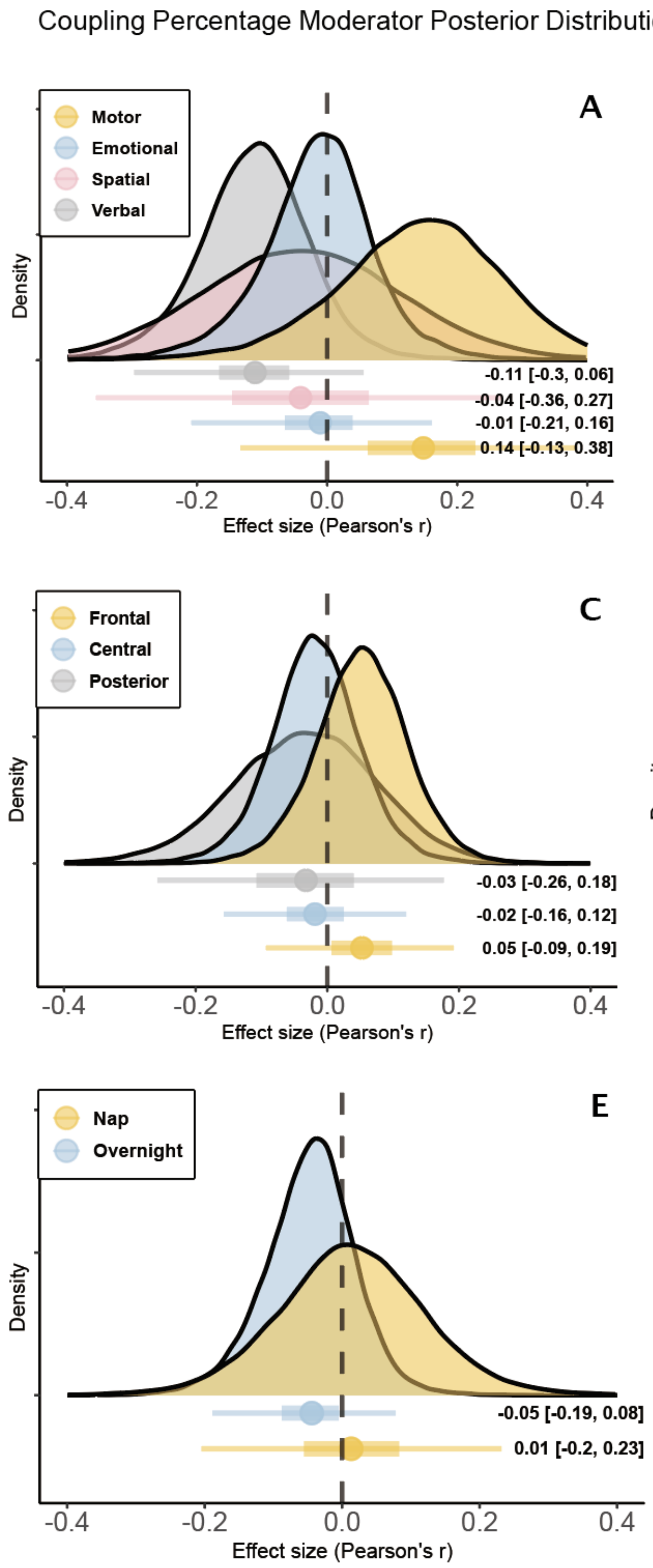


Figure S 8.4: Posterior distributions of moderation factors of the coupling percentage

Notes. The vertical line reflects the mean correlation coefficient under the null hypothesis. Colored dots and error bars reflect the mean and 95% credible intervals of each posterior distribution.

### 9 Interaction models and sensitivity analysis

Table S 9.1: Summary of interaction and sensitivity models for the coupling phase-memory association

| Models | Weight | Factors | Estimate (95% CrI) | Age/Year slope (95% CrI) |
|---|---|---|---|---|
| Age × Channel | 0.17 | Age × Frontal | 0.11 (-0.05, 0.27) | 0.001 (-0.005, 0.007) |
| | | Age × Central | 0.19 (0.04, 0.33) | -0.007 (-0.012, -0.001) |
| | | Age × Posterior | 0.12 (-0.18, 0.43) | -0.004 (-0.019, 0.012) |
| Age × Task | 0.06 | Age × Verbal | 0.10 (-0.05, 0.25) | -0.001 (-0.006, 0.004) |
| | | Age × Spatial | 0.16 (-0.21, 0.56) | -0.010 (-0.022, 0.002) |
| | | Age × Emotional | 0.22 (-0.38, 0.71) | -0.010 (-0.048, 0.028) |
| | | Age × Motor | 0.27 (-0.14, 0.68) | 0.009 (-0.028, 0.010) |
| Channel × Spindle | 0.16 | Frontal × Fast SP | 0.18 (0.06, 0.29) | |
| | | Central × Fast SP | 0.06 (-0.05, 0.16) | |
| | | Posterior × Fast SP | -0.01 (-0.18, 0.16) | |
| | | Frontal × Slow SP | 0.02 (-0.11, 0.14) | |
| | | Central × Slow SP | 0.04 (-0.09, 0.17) | |
| | | Posterior × Slow SP | -0.05 (-0.23, 0.14) | |
| Time-lag | 0.00 | Time-lag bias | | -0.010 (-0.050, 0.026) |
| Prior sensitivity | 0.02 | $N$(0,2.5), InvGamma(2, 0.5) | 0.07 (-0.01, 0.14) | |
| | 1.00 | Non-informative | 0.07 (0.01, 0.13) | |
| Main model | 1 - weight | None | 0.07 (0.01, 0.13) | |

Table S 9.2: Summary of interaction and sensitivity models for the SP amplitude-memory association

| Models | Weight | Factors | Estimate (95% CrI) | Age/Year slope (95% CrI) |
|---|---|---|---|---|
| Age × Channel | 0.00 | Age × Frontal | 0.17 (-0.02, 0.36) | -0.002 (-0.008, 0.003) |
| | | Age × Central | 0.15 (-0.05, 0.34) | -0.002 (-0.009, 0.004) |
| | | Age × Posterior | 0.13 (-0.28, 0.55) | -0.008 (-0.032, 0.016) |
| Age × Task | 0.73 | Age × Verbal | 0.10 (-0.09, 0.31) | 0.001 (-0.005, 0.007) |
| | | Age × Spatial | 0.38 (0.08, 0.67) | -0.009 (-0.016, -0.001) |
| | | Age × Emotional | 0.20 (-0.46, 0.73) | -0.015 (-0.058, 0.028) |
| | | Age × Motor | -0.26 (-0.80, 0.35) | 0.013 (-0.018, 0.044) |
| Channel × Spindle | 0.00 | Frontal × Fast SP | 0.14 (-0.03, 0.32) | |
| | | Central × Fast SP | 0.13 (-0.03, 0.28) | |
| | | Posterior × Fast SP | 0.05 (-0.17, 0.28) | |
| | | Frontal × Slow SP | 0.11 (-0.07, 0.28) | |
| | | Central × Slow SP | -0.01 (-0.20, 0.19) | |
| | | Posterior × Slow SP | -0.18 (-0.46, 0.10) | |
| Time-lag | 0.00 | Time-lag bias | | 0.000 (-0.070, 0.070) |
| Prior sensitivity | 0.00 | *N*(0,2.5), InvGamma(2, 0.5) | 0.07 (-0.04, 0.18) | |
| | 1.00 | Non-informative | 0.07 (-0.04, 0.19) | |
| Main model | 1 - weight | None | 0.07 (-0.04, 0.18) | |

Table S 9.3: Summary of interaction and sensitivity models for the coupling strength-memory association

| Models | Weight | Factors | Estimate (95% CrI) | Age/Year slope (95% CrI) |
|---|---|---|---|---|
| Age × Channel | 0.00 | Age × Frontal | 0.14 (-0.03, 0.30) | -0.001 (-0.007, 0.004) |
| | | Age × Central | 0.18 (0.01, 0.33) | -0.005 (-0.011, 0.001) |
| | | Age × Posterior | -0.06 (-0.38, 0.27) | 0.009 (-0.009, 0.026) |
| Age × Task | 0.00 | Age × Verbal | 0.11 (-0.04, 0.27) | -0.002 (-0.007, 0.003) |
| | | Age × Spatial | 0.13 (-0.18, 0.45) | -0.004 (-0.015, 0.007) |
| | | Age × Emotional | 0.17 (-0.39, 0.64) | -0.001 (-0.035, 0.035) |
| | | Age × Motor | 0.14 (-0.45, 0.67) | -0.004 (-0.028, 0.023) |
| Channel × Spindle | 0.00 | Frontal × Fast SP | 0.15 (0.01, 0.28) | |
| | | Central × Fast SP | 0.02 (-0.09, 0.14) | |
| | | Posterior × Fast SP | 0.08 (-0.12, 0.28) | |
| | | Frontal × Slow SP | 0.04 (-0.11, 0.18) | |
| | | Central × Slow SP | 0.05 (0.10, 0.20) | |
| | | Posterior × Slow SP | 0.15 (-0.05, 0.35) | |
| Time-lag | 0.43 | Time-lag bias | | 0.024 (-0.015, 0.060) |
| Prior sensitivity | 0.26 | $N$(0,2.5), InvGamma(2, 0.5) | 0.08 (0.00, 0.16) | |
| | 1.00 | Non-informative | 0.08 (0.02, 0.15) | |
| Main model | 1 - weight | None | 0.08 (0.02, 0.15) | |

Table S 9.4: Summary of interaction and sensitivity models for the coupling percentage-memory association

| Models | Weight | Factors | Estimate (95% CrI) | Age/Year slope (95% CrI) |
|---|---|---|---|---|
| Age × Channel | 0.40 | Age × Frontal | -0.04 (-0.76, 0.72) | 0.005 (-0.038, 0.045) |
| | | Age × Central | -0.00 (-0.59, 0.63) | -0.001 (-0.036, 0.031) |
| | | Age × Posterior | 0.18 (-0.61, 0.98) | -0.014 (-0.062, 0.030) |
| Age × Task | 0.65 | Age × Verbal | 0.42 (-0.24, 1.16) | -0.035 (-0.085, 0.008) |
| | | Age × Spatial | -0.04 (-1.00, 1.07) | 0.003 (-0.072, 0.072) |
| | | Age × Emotional | 0.21 (-0.54, 0.84) | -0.017 (-0.073, 0.034) |
| | | Age × Motor | 0.39 (-0.91, 1.49) | -0.015 (-0.085, 0.055) |
| Channel × Spindle | 0.00 | Frontal × Fast SP | 0.02 (-0.16, 0.19) | |
| | | Central × Fast SP | -0.04 (-0.20, 0.13) | |
| | | Posterior × Fast SP | 0.00 (-0.29, 0.27) | |
| | | Frontal × Slow SP | 0.10 (-0.10, 0.29) | |
| | | Central × Slow SP | 0.01 (-0.18, 0.19) | |
| | | Posterior × Slow SP | -0.07 (-0.36, 0.95) | |
| Time-lag | NaN | Time-lag bias | | Did not perform |
| Prior sensitivity | 0.09 | $N$(0,2.5), InvGamma(2, 0.5) | -0.04 (-0.17, 0.09) | |
| | 1.00 | Non-informative | -0.03 (-0.16, 0.07) | |
| Main model | 1 - weight | None | -0.03 (-0.15, 0.07) | |

### 10 Frequentist analysis

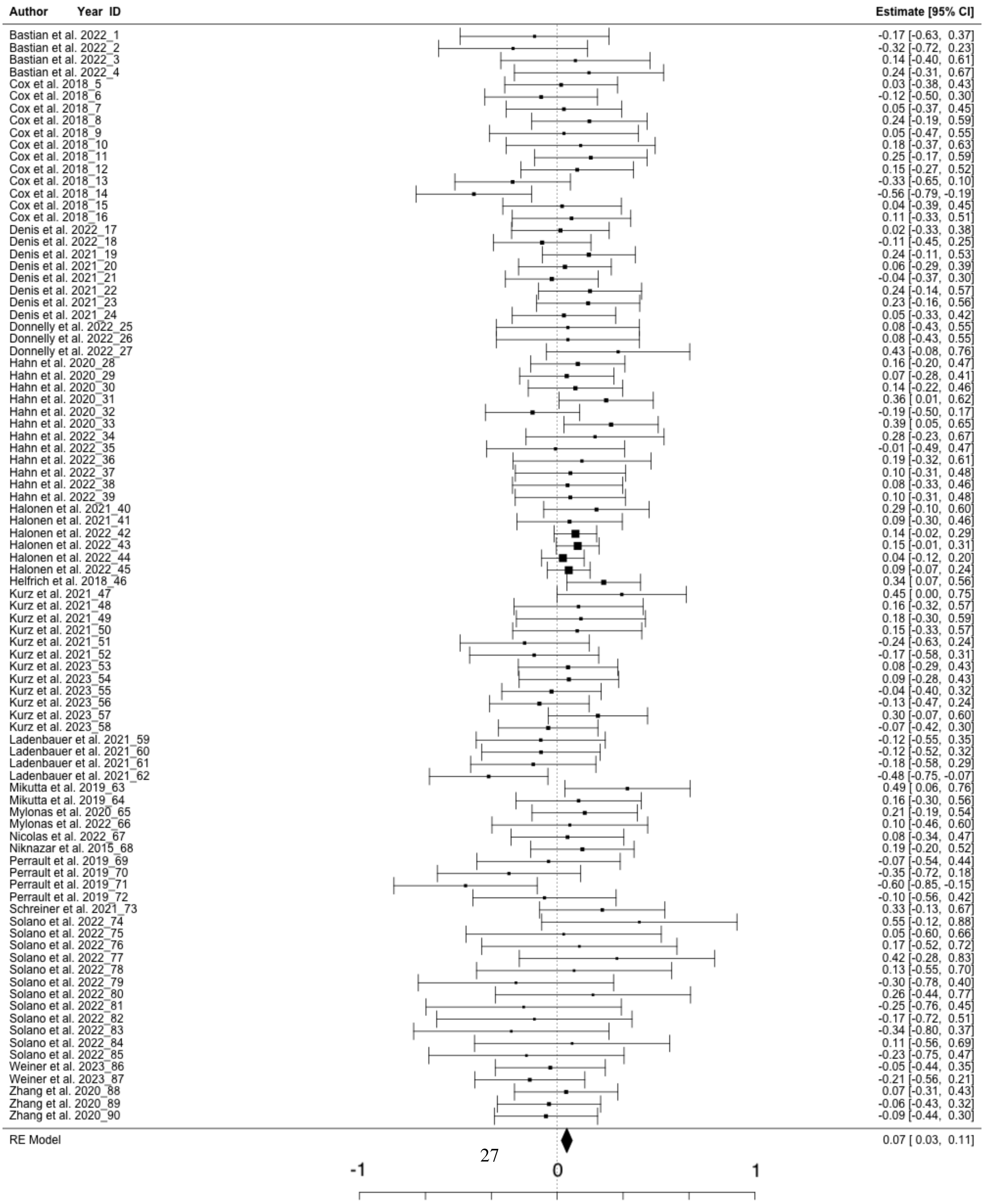


Figure S 10.1: Coupling phase frequentist forest plot

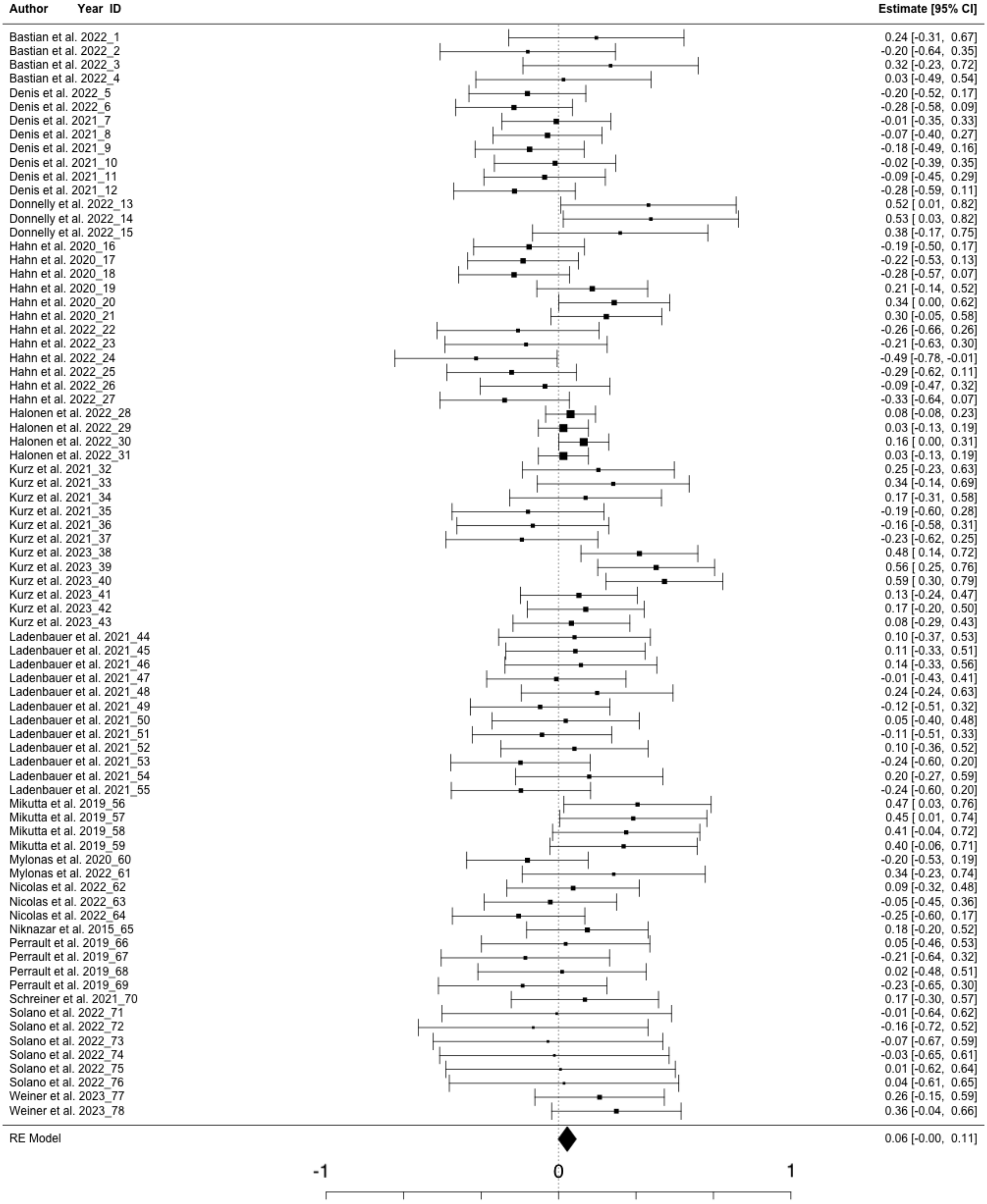




Figure S 10.2: SP amplitude frequentist forest plot

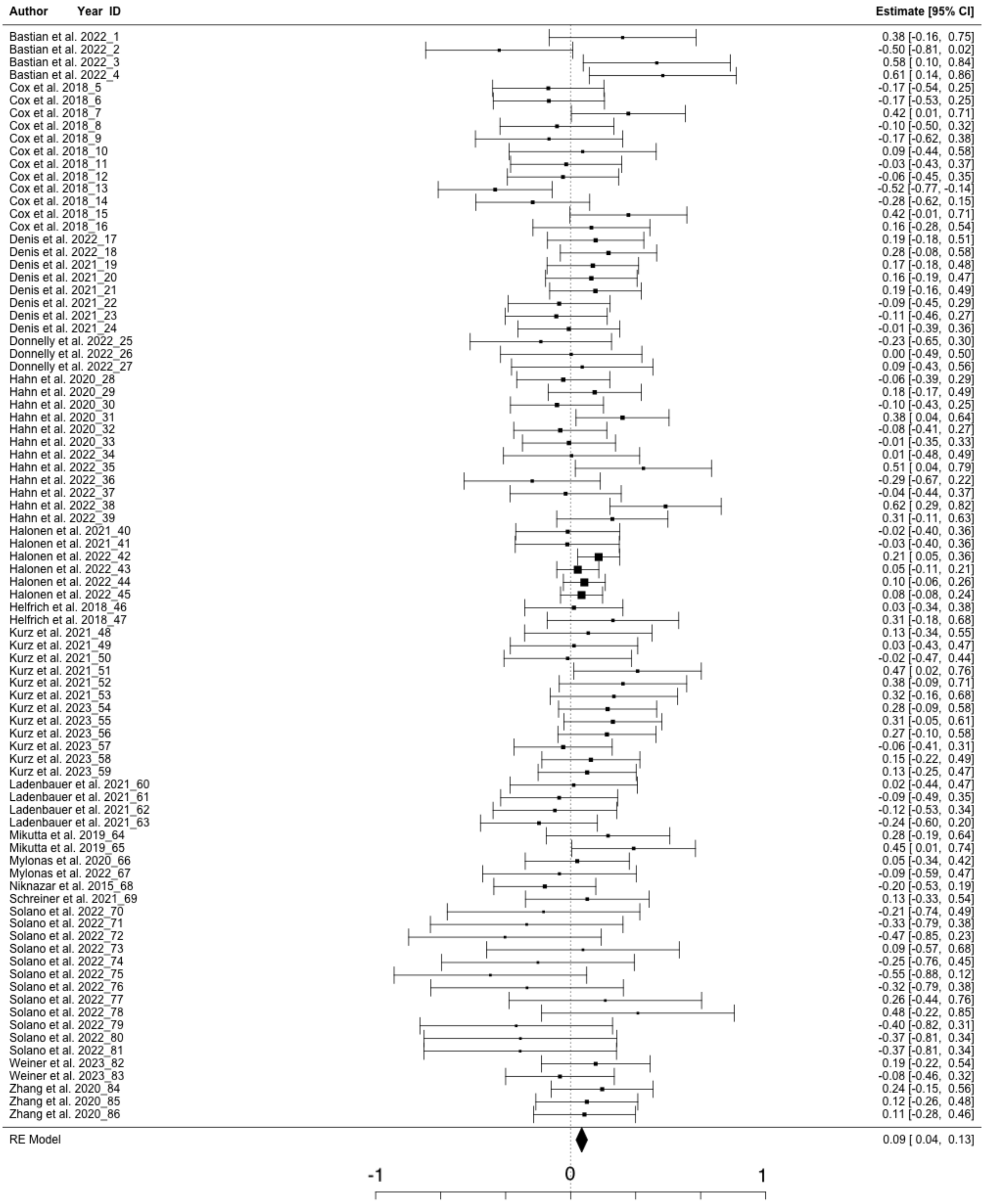




Figure S 10.3: Coupling strength frequentist forest plot

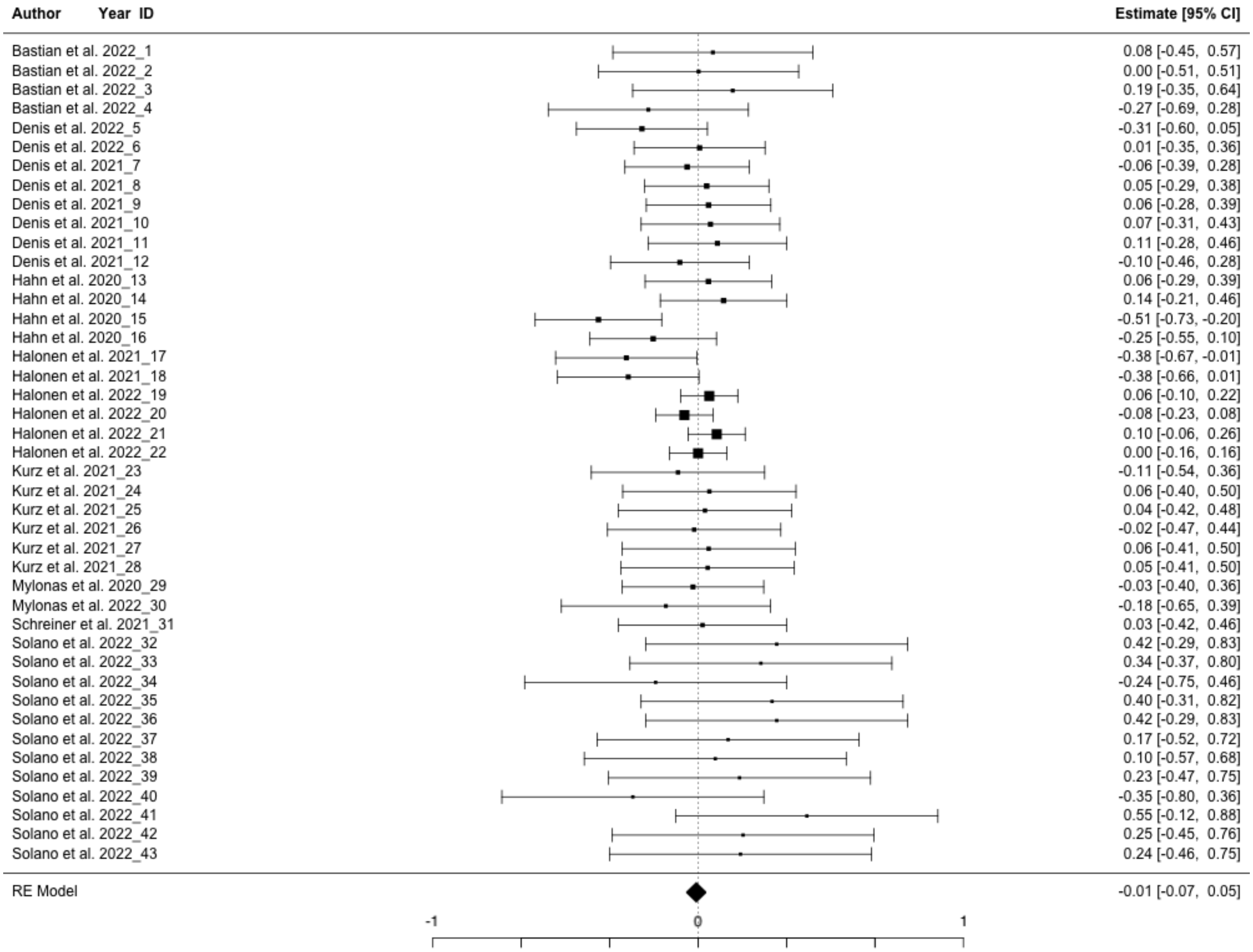


Figure S 10.4: Coupling percentage frequentist forest plot

### 11 Funnel plots for publication bias

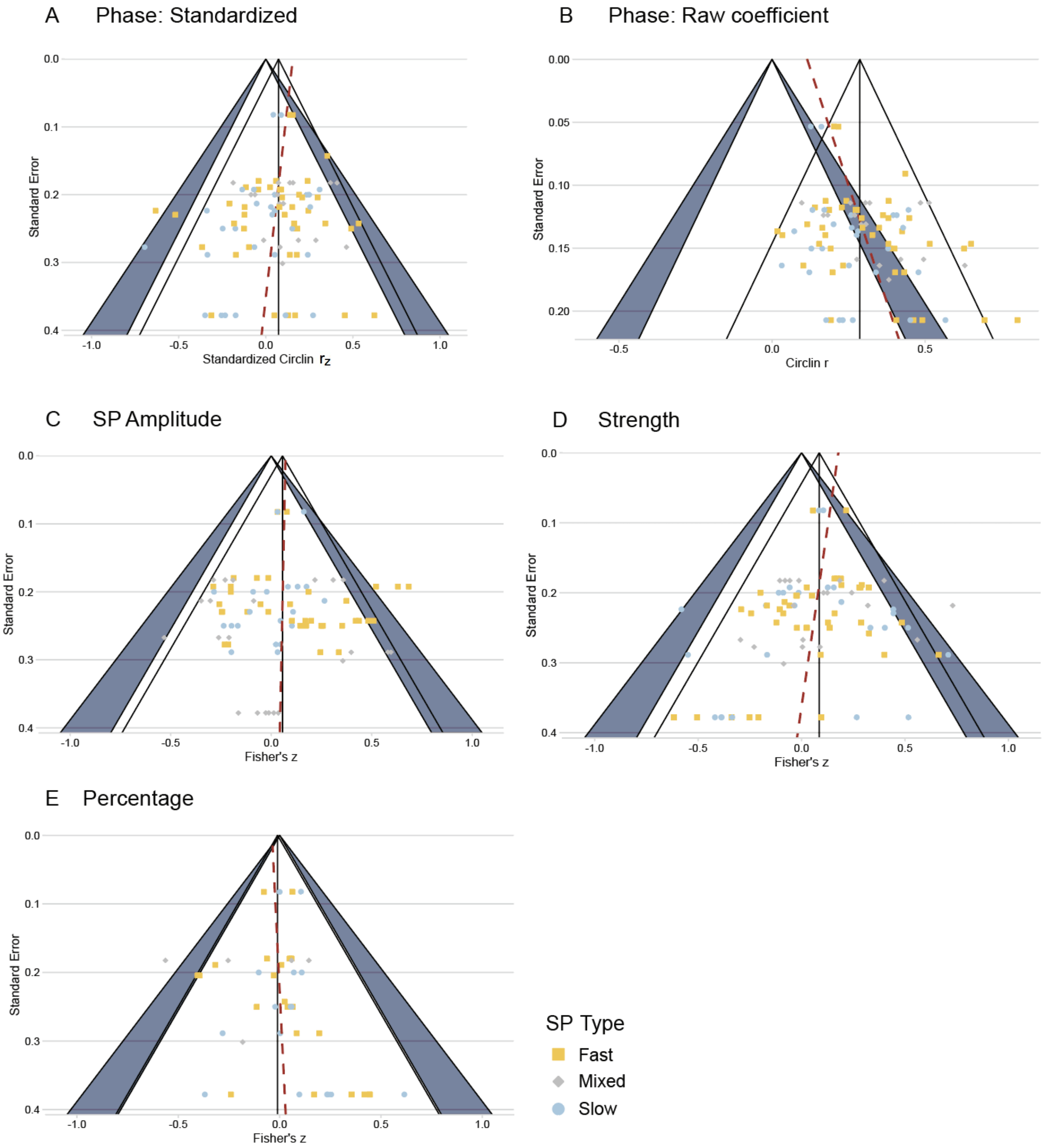


Figure S 11.1: Funnel Plot and Egger regression for assessing publication bias



Notes. Each colored dot represents one effect size and corresponding standard error. The outer border of transparent triangle indicates the area where it is anticipated that 95% of the included studies would fall if there were no publication biases present. The red dashed line represent the superimposed Egger's regression line. We noted that with sufficient sample sizes, the phase-memory association and strength-memory association tends towards a moderate magnitude.

#### 12 Simulation of circular-linear correlation and standardization

From Figure S12.1, we observed that the mean of the raw circular linear correlation coefficient (Circlin $r$) shifts with the change of sample sizes, and the distribution is highly skewed. By approximating the distribution of the weighted circular linear distribution to the chi-square distribution with 2 degrees of freedom[6], $\chi_2^2$, we derived that the circular-linear correlation coefficient has the following properties in an approximate form:

$$E(\rho) = \sqrt{\frac{\pi}{2n}},\ Var(\rho) = \frac{4-\pi}{2n},\ 0 \leq \rho \leq 1, n \geq 2$$

By standardizing the circular linear correlation coefficient (see Section 5.5.2), we first generated underlying populations that have null (0), moderate (0.3-0.4), or large (0.6-0.7) $r_z$ correlations, then tested whether the sampling distribution drawn from these populations followed a normal distribution across varying sample sizes.

We can observe from Figure S12.1 that under the null hypothesis, the standardized Circlin $r_z$ follows similar distributions with the linear Pearson's $r$. It is centered at 0 and can approximate unbounded normal after Fisher's $z$ transformation. The difference between Pearson's $r$ and standardized Circlin $r_z$ is that the negative x-axis of the circlin $r_z$ represents a magnitude of correlation under the null hypothesis without effects, instead of a negative direction of the association.

As long as the sample size is more significant than 25, the performance of the Circlin transformation is highly stable. However, we should acknowledge that the non-linear correlation is unreliable under small sample sizes. Therefore, we do not recommend conducting circular-linear correlational analysis when $n < 15$.

To ensure the normality of standardized Circlin $r_z$ under large effect size of correlations, we observed that sampling distributions when $r_z$ = 0.3-0.4 and 0.6-0.7 (Figure S12.2, 12.3) after Fisher's z transformation consistency fit superimposed normal distributions, and even perform robustly in conditions where sample sizes are relatively small.

Therefore, we encourage future studies to report the standardized coefficient instead of the raw Circlin $r$, which can accurately reflect the true effect size without exaggeration when the sample size is small, as well as improve the clarity of interpretation and comparability with other types of correlation coefficients across studies.

The code used in R to standardize the Circlin coefficient is: limma::zscore(n*r$^2$, dist='chisq', where, df=2)/sqrt(n), where $n$ represents the sample size while $r$ represents the Circlin coefficient.

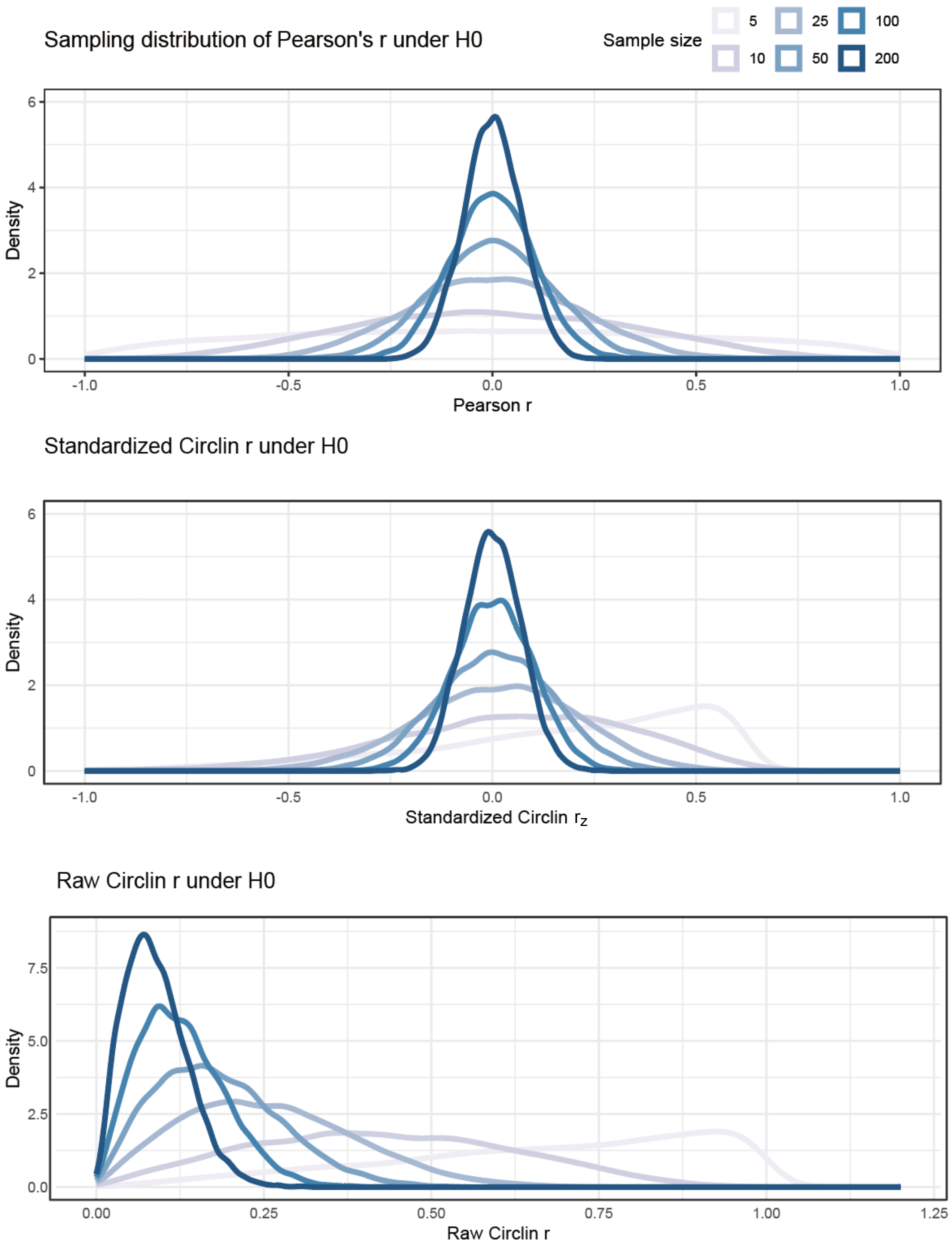


Figure S 12.1: Comparison of sampling distributions of standardized Circlin $r_z$ under null

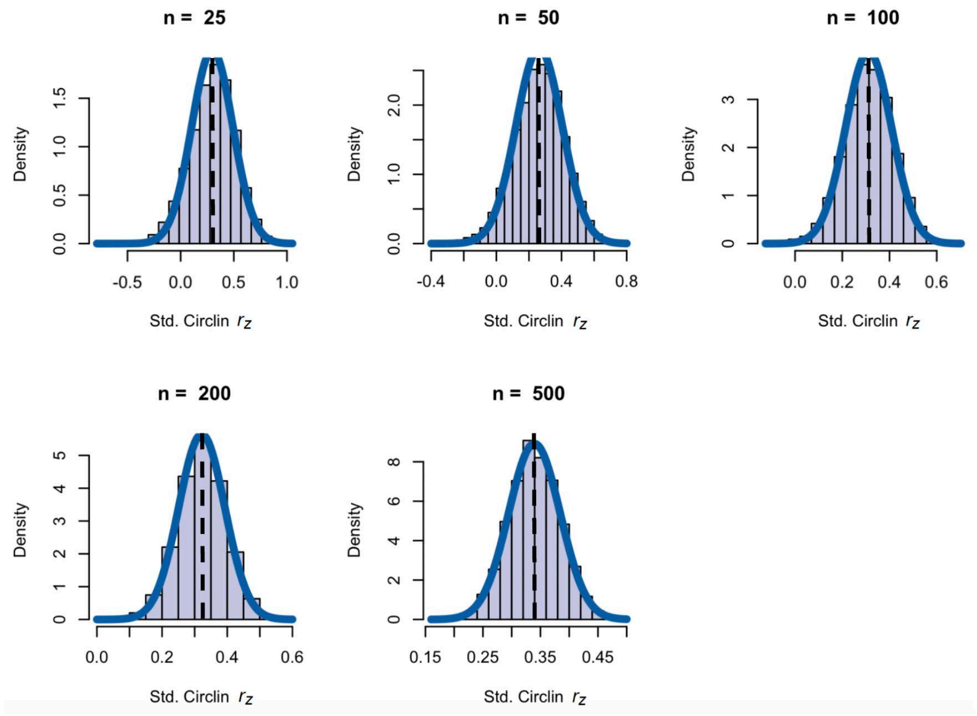


Figure S 12.2: Sampling distributions of standardized Circlin $r_z$ drawn from moderate correlations ($r_z$ = 0.3-0.4). Dashed vertical lines represent the true $r_z$ of the population used to draw samples.

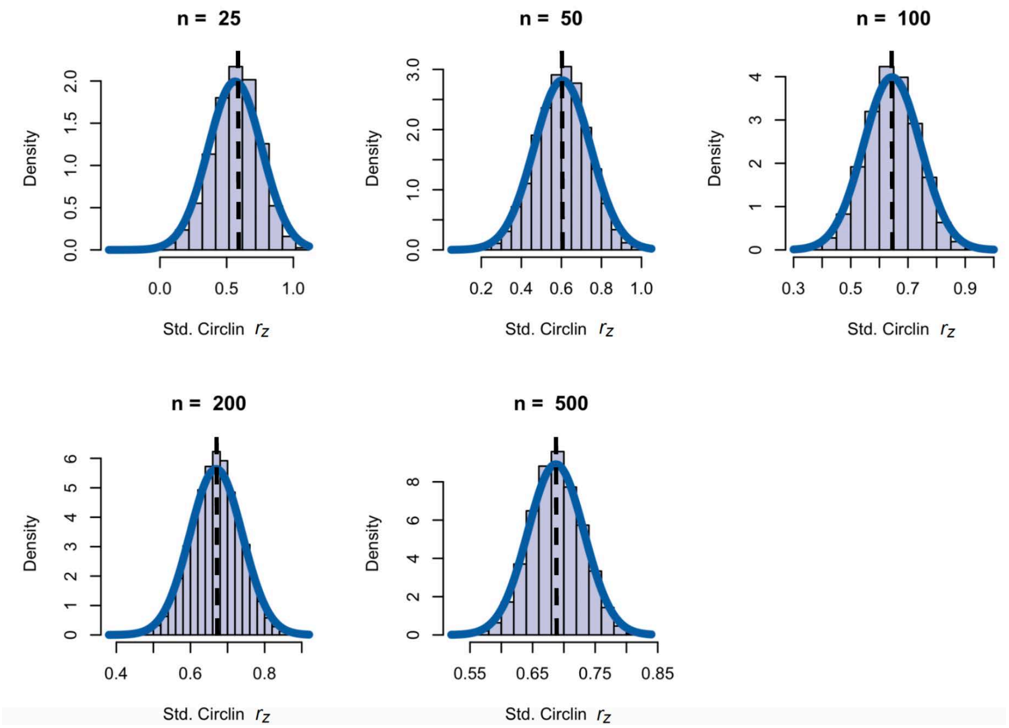


Figure S 12.3: Sampling distributions of standardized Circlin $r_z$ drawn from strong correlations ($r_z$ = 0.6-0.7). Dashed vertical lines represent the true $r_z$ of the population used to draw samples.

### 13 SO-slow SP spatial-temporal analysis

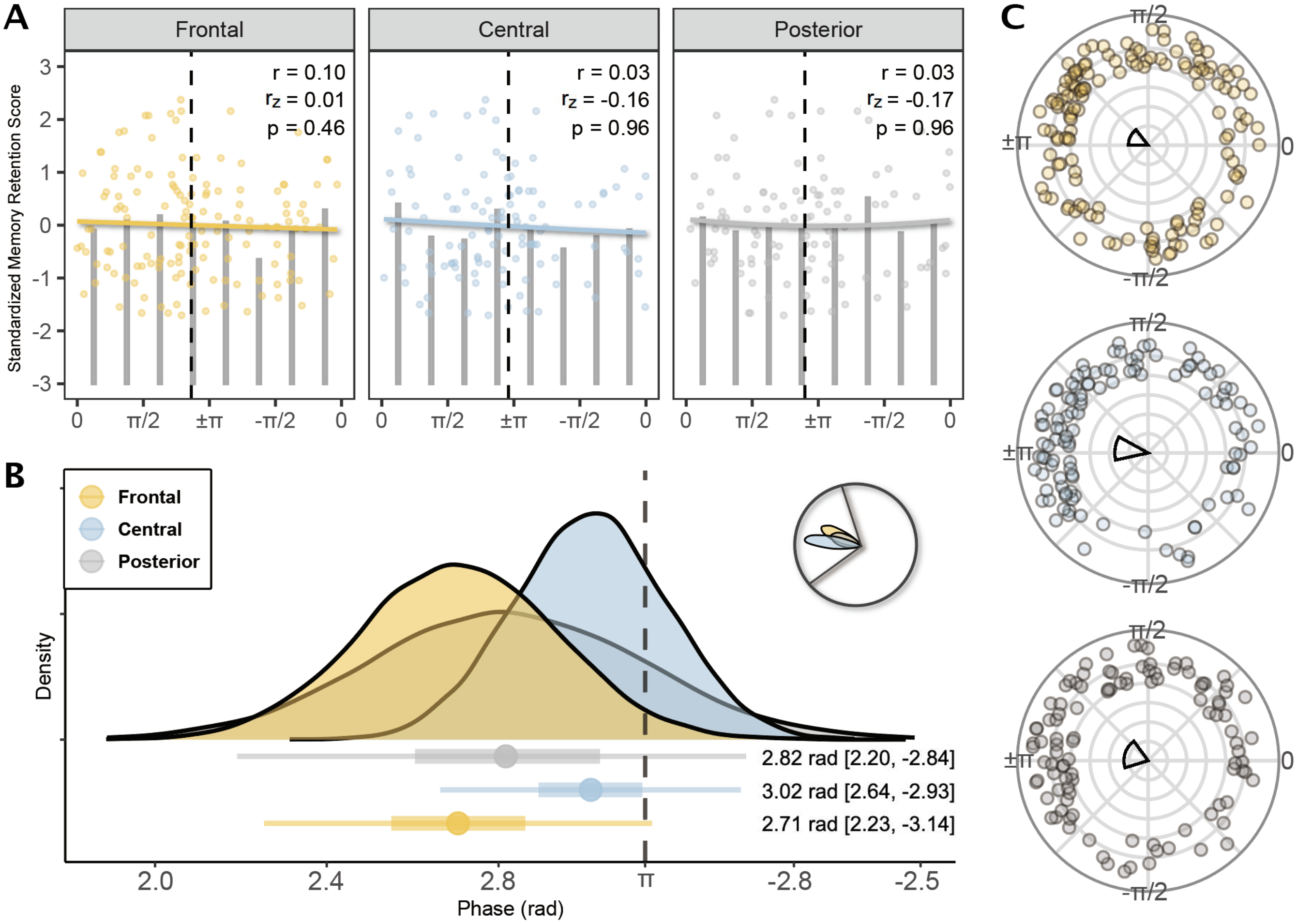


Figure S 13.1: Preferred slow oscillation-slow spindle coupling phase and its association with the memory retention

Notes. (A) Quadratic regression of the phase-memory association under different regions of PSG channels aggregated from studies included in the meta-analysis. 0° peak of SO upstate; ±$\pi$° trough of SO downstate; $r$ circular-linear correlation coefficient; $r_z$ standardized circular-linear correlation coefficient. Bars represent the mean memory retention scores per $\pi$/4 radian (45°). Dashed vertical line represents the mean preferred phase across studies. Colored quadratic fit line represents the direction of the relationship. None of the PSG channels displays a typical quadratic relationship around the down-state trough of SOs. (B) Posterior distributions of mean preferred phases from the Bayesian circular mixed-effect model. The circular posterior distribution has been reported in the top-right corner, and the area between two black lines has been projected in a linear scale in the main graph. The vertical line reflects the down-state trough of SOs. Dots and error bars reflect the mean and 95% credible intervals of phases detected from each channel c luster. Phase values have been reported as degrees. (C) Circular plot of the preferred coupling phase. Arranged from top to bottom in the order of frontal, central, and posterior. The direction of each colored dot represents the preferred coupling phase of each subject recorded from PSG channels in each cluster. The direction of the mean resultant vector indicates the mean preferred coupling phase across subjects, the width indicates the 95% credible interval of the mean coupling phase, while the length from 0 (center) to 1 (circumference) indicates the consistency of coupling phase across subjects.

Contrary to fast SPs, we found that the preferred phase of SO-slow SP coupling did not show significant quadratic association with memory retention in any topographic regions (see Figure S13.1A), all $r \leq 0.1$, $r_z \leq 0.01$, $p > 0.05$. After taking into account the repeated measurement, we found that SO-slow SP coupling happens slightly before the down-state trough of SOs, reflected by the phase in frontal (2.71 rad [2.23, -3.14]), central (3.02 rad [2.64, -2.93]) and posterior regions (2.82 rad [2.20, -2.84]). However, the phase distribution across participants is considerably less consistent than the SO-fast SP coupling (all $z < 0.34$, $p < 0.01$, Rayleigh test), which might be relevant to between-study variabilities on the definition of slow SPs.

No significant phase shift has been found between frontal and central areas, $r = \Delta$ -0.31 rad, $BF_{10}$ = 0.06, probability = 0.05; between frontal and posterior areas, $r = \Delta$ -0.11 rad, $BF_{10}$ = 0.49, probability = 0.33; or between central and posterior areas, $r = \Delta$ 0.20 rad, $BF_{10}$ = 3.64, probability = 0.78. In summary, there is no strong evidence to support the association between the SO-slow SP coupling phase and memory retention performance, or to substantiate phase shifts across cortical areas.
